# A pangenomic approach identifies new structural variants associated with gene expression in the CARTaGENE cohort

**DOI:** 10.64898/2026.11.24.768960

**Authors:** Rose Laflamme, Cristian Groza, Guillaume Bourque, Guillaume Lettre

## Abstract

Structural variants (SVs) are major contributors to the genetic diversity of human populations. While many small variants have been associated with human traits and diseases, the challenge to genotype SVs using short-read DNA sequencing and a linear reference genome has made association testing with SVs more difficult. To address these limitations and investigate the associations of SVs with complex human traits, we used PanGenie, a pangenome-based short-read genotyper, together with the second release of the Human Pangenome Reference Consortium (HPRC) pangenome to genotype 645,680 SVs in 2,173 participants from the CARTaGENE cohort. Through conditional eQTL analyses that account for associations attributable to known variation identified using a standard approach based on the GRCh38 reference genome, we identified 103 independent SV-eGene associations involving 99 unique SVs (SV-eQTLs). Of these associations, 41 replicated in the 1000 Genomes Project cohort (same direction of effect, nominal P-value <0.05). The 99 SV-eQTLs were more likely than expected to overlap with candidate cis-regulatory elements as defined by the ENCODE Project (P-value <0.002). Finally, we found that 65/99 (66%) of the SV-eQTLs were in strong linkage disequilibrium (*r*^2^>0.8) with small genetic variants (SNPs and indels) that belong to Open Targets fine-mapping credible sets, suggesting that these SVs are candidate causal variants for associations identified by genome-wide association studies. Our results highlight new opportunities offered by pangenome methods to more comprehensively integrate SVs into the study of human phenotypic variation.

## BACKGROUND

Structural variants (SVs) such as deletions, insertions and duplications are defined as large genomic alterations encompassing at least 50 DNA base pairs (bp) (1–3). Because they span a wide range of sizes and types, SVs are considered the largest contributors to genetic diversity within populations (1,2,4,5). As with small variants (single nucleotide polymorphisms [SNPs] and insertions-deletions [indels]), SVs have been associated with many human phenotypes and diseases (2,6,7). However, most genome-wide association studies (GWAS) have focused on small variants because they are easier to genotype (4,7). SV detection is possible from short-read whole-genome sequencing data (SR-WGS), but more than two-thirds remain uncaptured (3,8,9). This limitation arises because short reads cannot fully span large SVs or because reads containing SVs may misalign when the corresponding sequence is absent from the reference genome (e.g. GRCh38) (1,4,10–12).

Long-read whole-genome sequencing (LR-WGS) has emerged as a powerful approach for SV discovery (13–17). By generating reads several kilobases long, LR-WGS can cover larger genomic regions, including repetitive regions, thus enabling the detection of larger and more complex variants with improved accuracy (13,16,17). High-quality haplotype-resolved genome assemblies derived from LR-WGS have led to the development of SV imputation panels (14,15), as well as pangenomic references for SV genotyping (9,16,18–22). A pangenome consists of a collection of genome sequences represented as a graph structure that captures genetic variation across multiple individuals (11,23). Unlike the current reference genome (i.e. GRCh38), which represents a single haplotype, pangenome graphs allow multiple alleles at variable sites, including SVs (11,23). Many research groups have generated pangenome references to overcome the limitations of a single linear reference genome and to better represent the genetic diversity of their populations of interest (9,12,16,18–21). For instance, the Human Pangenome Reference Consortium (HPRC) released a first draft human pangenome reference (HPRC1) in May 2023, built from 47 diploid assemblies (9). Their second draft pangenome (HPRC2), released in May 2025, was built from 460 diploid assemblies (16).

To investigate the association between SVs and complex human phenotypes, we used data from the CARTaGENE cohort (CaG). CaG is the largest population-based cohort in the province of Quebec (Canada) (24,25). It includes 43,032 participants aged between 40 and 69 years at baseline recruited in different metropolitan regions of Quebec (24,25). High-coverage SR-WGS data is available for 2,173 CaG participants, including 1,756 participants with French-Canadian ancestry (QFC), 163 with four grandparents born in Haiti (QHA), and 131 with four grandparents born in Morocco (QMO) (24). Haiti and Morocco are both important sources of immigration in Quebec, but most of the Quebec population is QFC, descending from French settlers who colonized the territory in the 17^th^ and 18^th^ centuries (24).

Here, we used a pangenomic approach to characterize new SVs in CaG and test their association with gene expression. We used the graph genomes from HPRC2 and the software PanGenie to genotype SVs in 2,173 CaG participants. In 504 CaG participants with whole-blood RNA-sequencing (RNAseq) data available, we applied conditional analyses to discover novel associations with gene expression (i.e. expression quantitative trait loci or eQTL) that are solely driven by SVs identified through our pangenome strategy. We annotated these new eQTLs using regulatory elements predicted by ENCODE and GWAS fine-mapping results from Open Targets (26–28). Our results indicate that pangenome-augmented genetic datasets can identify novel associations with complex human phenotypes.

## RESULTS

### Genotyping SVs from HPRC

We used the algorithm PanGenie (29), a pangenome-based short-read genotyper, to genotype genetic variation from HPRC1 and HPRC2 in 2,173 CaG participants for whom SR-WGS data was available. We used the Minigraph-Cactus HPRC graphs with CHM13 as the reference sequence. We genotyped 318,162 SVs from HPRC1 with PanGenie (referred to as *HPRC1+PanGenie*: 68,161 deletions, 171,703 insertions, and 77,298 complex SVs) and 645,680 SVs from HPRC2 (referred to as *HPRC2+PanGenie*: 103,630 deletions, 455,071 insertions, and 86,979 complex SVs) (**Supplementary Figure 1**). We defined complex SVs as variants for which both the reference and alternate alleles are longer than 1 bp, with at least one allele neasuring ≥50 bp. Note that PanGenie does not distinguish duplications from insertions, and that multiallelic variants are represented as multiple biallelic variants in the final datasets (**Methods**). Using standard SR-WGS-based SV detection methods (i.e. based on the GRCh38 reference genome, **Methods**), we previously genotyped 25,820 SVs in the same 2,173 CaG participants, including 24,022 deletions and 1,798 duplications (this SR-WGS-only approach did not identify insertions nor complex SVs [**Supplementary Figure 1**]) (24). This is referred as the S*tandard SV* dataset in subsequent sections.

We observed that only a small subset of the SVs identified in the *Standard SV* dataset were also genotyped using the HPRC-based approach. For instance, if we focus on deletions that overlap perfectly (same breakpoint coordinates), only 2,281 deletions were found in both the *Standard SV* and *HPRC1+PanGenie* datasets (2,508 deletions in *Standard SV* and *HPRC2+PanGenie*). When using a 50% reciprocal overlap to match deletions across datasets, there were 6,823 and 9,709 deletions in the *HPRC1+PanGenie* and *HPRC2+PanGenie* datasets, respectively, that overlap with deletions in the *Standard SV* dataset. A possible explanation for these differences is that many SVs might be specific to (or enriched in) the CaG populations that were sequenced (QFC, QHA, QMO); these variants would not be present in HPRC and therefore not genotyped by PanGenie. Accordingly, although there are less SVs in the *Standard SV* dataset when compared to the HPRC-based datasets, the proportion of SVs with alternate allele frequency <0.1% in the *Standard SV* dataset is higher, suggesting population specificity for rarer SVs (**Supplementary Figure 2**).

### Assessing SV genotype concordance using LR-WGS

To evaluate SV genotype accuracy, we used LR-WGS data from high-coverage PacBio HiFi sequencing available for 20 QFC participants generated as part of a different project (**Methods**) (30). In the *HPRC2+PanGenie* dataset, after matching SVs based on identical reference and alternate alleles, we found 28,595 SVs available in the LR-WGS dataset, including 11,081 deletions, 17,038 insertions and 476 complex SVs. Overall, 92.7% of the genotypes were concordant between the LR-WGS and the *HPRC2+PanGenie* call sets, with concordance rates of 93.5% for deletions, 92.1% for insertions and 93.5% for complex SVs. The overall concordance rate between the LR-WGS and *HPRC1+PanGenie* datasets was 92.2% (94.01% for deletions, 90.5% for insertions, and 94.6% for complex SVs). These genotype concordance rates are comparable to the accuracy reported previously for PanGenie (29). As a comparison, we calculated a genotype concordance rate of 98.5% for 850 deletions in the *Standard SV* dataset when compared to the LR-WGS genotypes. We hypothesize that the higher genotype concordance for the *Standard SV* dataset can be explained by the quality-control filters applied with the standard approach. Indeed, the standard approach retained only SVs detected by two SV callers per sample, creating a confident subset of true positives. In other words, the SR-WGS-based standard approach only discovered a subset of “simple” SVs in regions where short reads perforn well, whereas the pangenome includes more complex SVs in regions that are difficult to genotype.

### SV diversity across ancestral groups

We quantified the number of SVs genotyped per ancestral group (QFC, QHA and QMO) by counting non-monomorphic SVs. Using *HPRC1+PanGenie*, 54.2% of SVs were shared among all three groups, while 7.2%, 14.1% and 0.8% were unique to the QFC, QHA and QMO groups, respectively (**Figure 1A**). Because we used a pangenome-based genotyping approach, our analyses were limited to the genetic variation represented in the reference graphs. Consequently, populations more similar to those represented in the reference pangenome may have better representation and are expected to have a greater number of successfully genotyped SVs. Since 51% of the assemblies in HPRC1 are from African-ancestry participants and only 2% from European-ancestry individuals, this likely explains why QHA, which is genetically closer to the African samples, had twice as many unique SVs as the QFC ancestry, which is closer to the European samples, despite having nearly tenfold fewer samples (N_QFC_=1,756 vs N_QHA_=163) (9). Using *HPRC2+PanGenie*, 35.3% of SVs were shared among all three groups, while 18.0%, 17.4% and 1.7% were unique to QFC, QHA and QMO, respectively (**Figure 1B**). Again, the comparable number of SVs unique to the QFC and the QHA groups may be explained by the populations represented in HPRC2 (13% for European-ancestry and 29% for African-ancestry). As expected, 35% of SVs in the *Standard SV* dataset were uniquely identified in QFC, reflecting the fact that this group represented the majority (81%) of the CaG SR-WGS data (**Figure 1C**).

**Figure 1.**
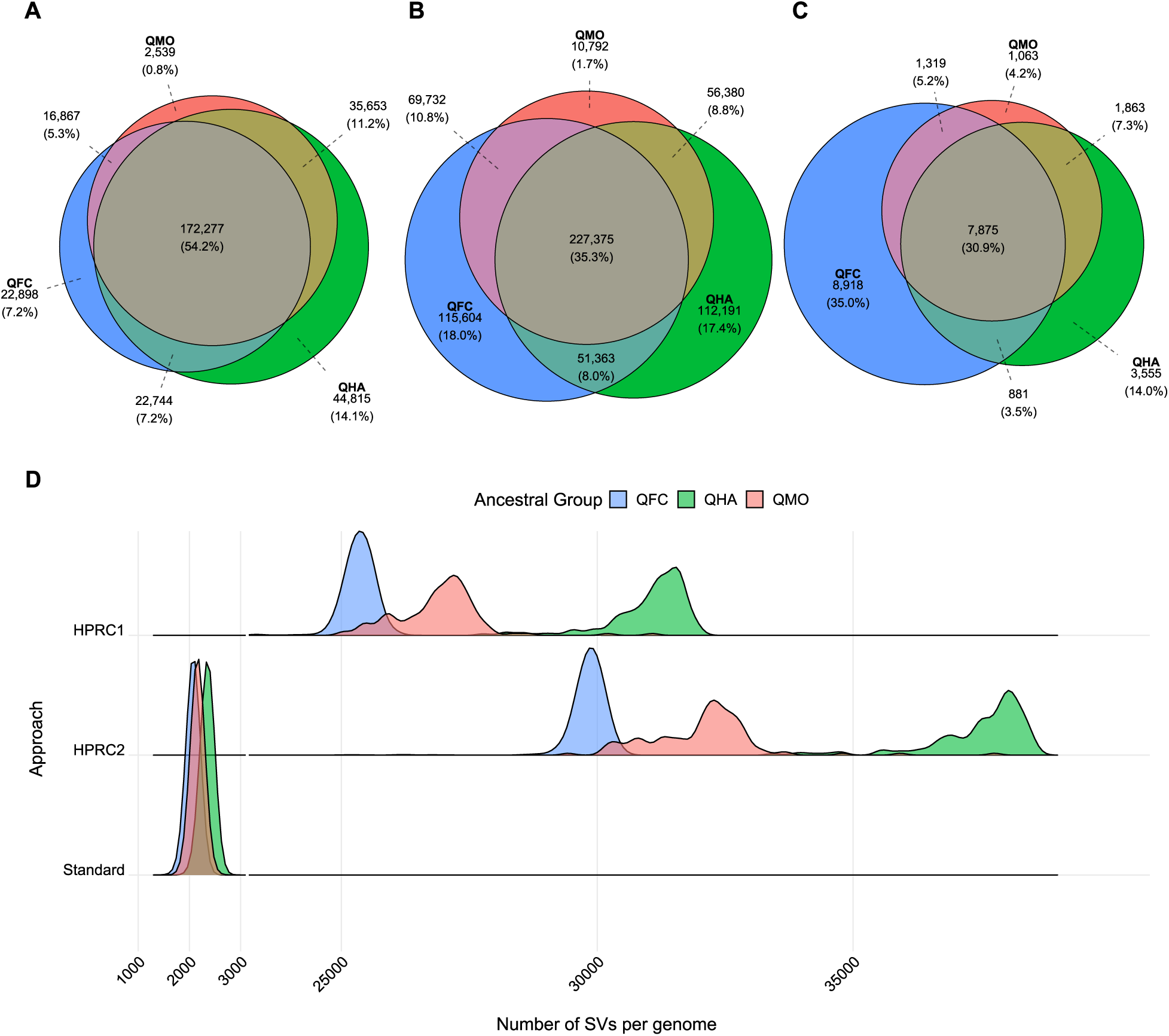
SV representation across CaG ancestral groups. Number of SVs genotyped in 2,173 CaG participants per ancestral group using (**A**) *HPRC1+PanGenie*, (**B**) *HPRC2+PanGenie* or (**C**) the *Standard SV* detection approach based on the GRCh38 reference genome. The ancestral groups are Quebec residents of French-Canadian ancestry (QFC, blue circles, N=1,756), Quebec residents of Haitian ancestry (QHA, green circles, N=163) and Quebec residents of Moroccan ancestry (QMO, red circles, N=131). The number of SVs genotyped per ancestral group is the number of SVs with a minor allele frequency >0 in the CaG participants from each ancestral group. (**D**) Number of SVs genotyped per genome in 2,173 CaG participants using different approaches: *HPRC1+PanGenie* (HPRC1), *HPRC2+PanGenie* (HPRC2), and *Standard SV* detection approach based on the GRCh38 reference genome (Standard). The x-axis represents the number of SVs genotyped per genome. The y-axis represents the relative density.

We also quantified the number of SVs genotyped per genome for each ancestral group. We counted an SV if a participant carried at least one non-reference allele. Compared with the other ancestral groups, QHA showed the highest number of SVs per genome, consistent with the increased genetic diversity observed in populations of African ancestry (31) (**Figure 1D**). QFC showed the fewest number of SVs per genome, with a mean of 25,338, 29,822 and 2,071 SVs identified using *HPRC1+PanGenie*, *HPRC2+PanGenie* and the *Standard SV* approach, respectively (**Figure 1D**).

### eQTL discovery

Given that HPRC2 allowed us to genotype more SVs in the CaG samples, we used SV calls from the *HPRC2+PanGenie* dataset for downstream analyses. We performed cis-eQTL analyses to test the association between expressed genes (eGenes) and SV genotypes using tensorQTL (32). We tested the association between 161,624 common SVs (minor allele frequency (MAF) ≥1%) and the expression of 13,558 genes in 504 CaG participants with whole-blood RNAseq data available (N_QFC_=471, N_QHA_=5, N_QMO_=3 and 25 additional CaG participants of European ancestry). We identified 11,266 genome-wide significant cis-eQTL at P-value<5×10^-8^ (**Supplementary Figure 3A**). In contrast, when using the *Standard SV* dataset on the same samples, we tested 12,991 genes and 4,653 SVs, and found 720 significant cis-eQTL (**Supplementary Figure 3B**).

Many of the cis-eQTL associations described above implicate SVs that are in linkage disequilibrium (LD) with other variants, including SNPs, indels but also SVs identified using the *Standard SV* SR-WGS-based method. To evaluate the added value of our pangenomic approach, we developed a pipeline to identify eQTL signals driven by *HPRC2+PanGenie* SVs. For each gene, we identified independent cis-eQTL signals explained by SNPs, indels and *Standard* SVs, and included the genotypes of these independent variants as covariates in models to test the association between gene expression and *HPRC2+PanGenie* SV genotypes. These conditional cis-eQTL analyses identified 186 genome-wide significant SV-eGene pairs (at P-value <5×10^-8^), corresponding to 171 unique SVs and 101 unique eGenes (**Figure 2A**). We further conditioned on *HPRC2+PanGenie* SVs to ensure that these cis-eQTL signals were independent from each other. This strategy highlighted 103 independent SV-eGene pairs (99 unique SVs) (**Table 1** and **Supplementary Table 1**), including one additional genome-wide significant association with the expression of *B3GNTL/ǪTGAL* (**Figure 2B-C**). These 99 SV-eQTLs included 44 deletions, 49 insertions and 6 complex SVs. Interestingly, 71/103 (69%) SV-eGenes pairs were not genome-wide significant before conditional analyses on variants detected using the *Standard* approach (**Supplementary Table 1** and **Supplementary Figure 4**). The 99 SV-eQTLs were more common (median alternate allele frequency 18.7% vs 3.9% and mean alternate allele frequency 22% vs 11%) and had higher genotype quality than the 161,624 *HPRC2-PanGenie* SVs tested in the eQTL analyses (**Supplementary Figures 5-6**). Finally, the majority 68/99 (68%) of the SV-eQTLs identified by our pangenomic approach are not available in the ImputeSV reference panel and would have therefore not been discovered by SV imputation (14).

**Figure 2.**
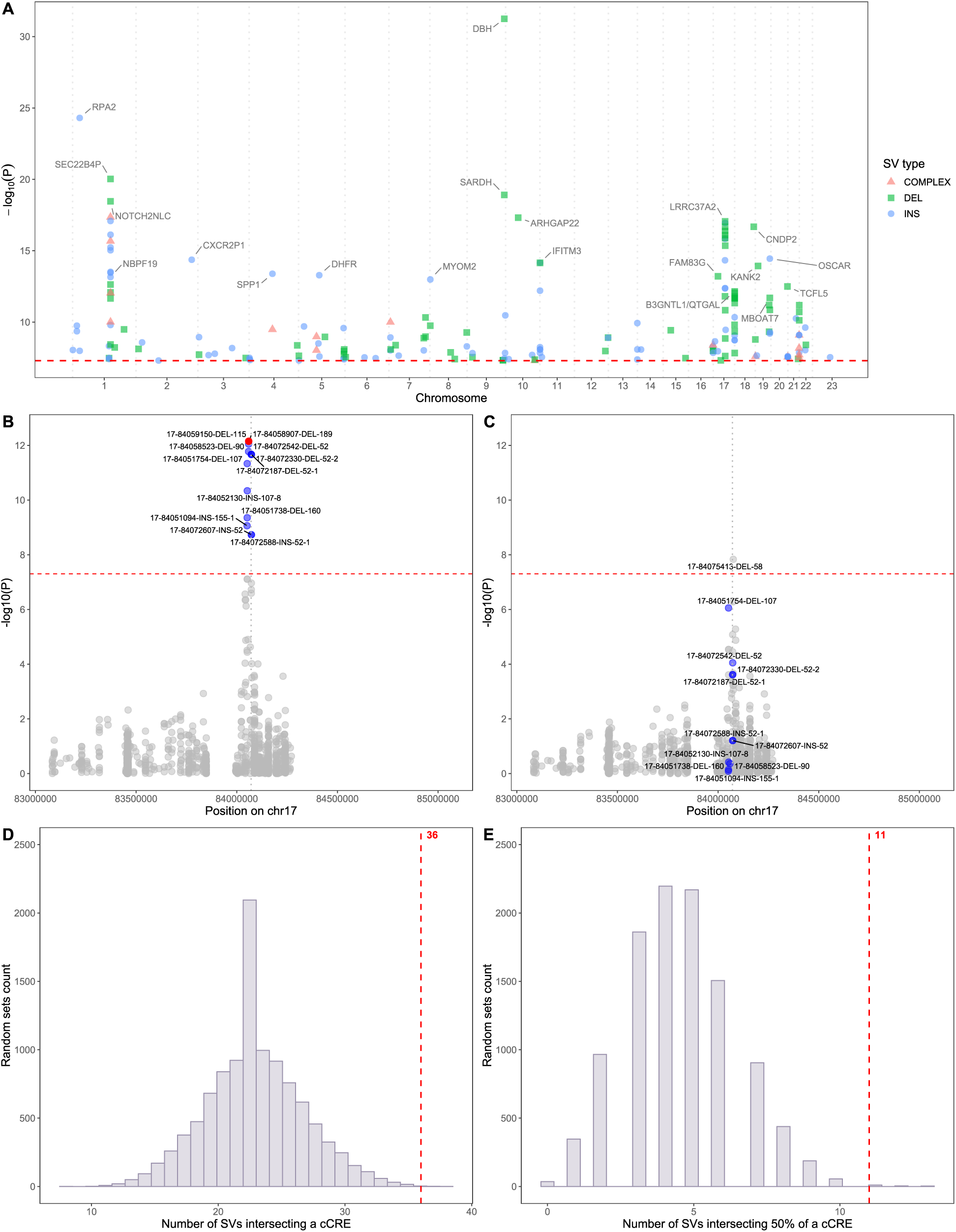
Conditional analyses identify pangenome-based SVs associated with gene expression. (**A**) Manhattan plot of the association results between SVs from the *HPRC2+PanGenie* dataset and gene expression in 504 CaG participants. Shown are SVs that remain significant after conditioning on SNPs, indels and SVs from the *Standard SV* dataset. The red line represents the threshold used to denote genome-wide significance for cis-eQTL (P-value=5×10^-8^). 186 SV-eGenes pairs were identified, corresponding to 99 unique and independent SVs. (**B**) LocusZoom plot of the association results between *HPRC2+Pangenie* SVs and the expression of *B3GNTL/ǪTGAL*. Results are conditioned on genotypes at nearby SNPs, indels and *Standard* SVs. The red dot is the most significant *HPRC2+Pangenie* SV at the locus whereas blue dots correspond to SV with P-value <5×10^-8^ (red dashed line). The vertical dotted line corresponds to the transcription start site of *B3GNTL/ǪTGAL*. (**C**) Identification of an additional *HPRC2+PanGenie* SV associated with *B3GNTL/ǪTGAL* expression levels when we conditioned on nearby SNPs, indels and *Standard* SVs, as well as the top *HPRC2+PanGenie* SV at the locus (17-64059150-DEL-115, a 115 kb deletion, red dot in **B**). (**D**-**E**) Enrichment of SV-eQTLs that (**D**) minimally overlap (³1 bp) or (**E**) partially overlap (50%) ENCODE cCREs. The vertical red dotted lines indicate the number of SV-eQTLs that overlap cCREs: 36/93 (39%) for minimal and 11/93 (12%) for minimal and partial overlaps, respectively. The grey bars correspond to the distribution of overlapping SVs-cCREs among 10,000 sets of SVs matched based on SV type and cumulative length.

**Table 1.** SVs from the *HPRC2+PanGenie* dataset that are independently associated with gene expression. POS, position of the SV on build CHM13. LENGHT, length of the SV in base pairs. eGene, gene for which the expression is associated with the SV in the conditional eQTL analysis. P-value, P-value in the conditional eQTL analysis. Beta, effect size (in standard deviation units) in the conditional eQTL analysis. cCRE(rE2G), number of ENCODE candidate cis-regulatory elements intersected by the SV (number of ENCODE candidate cis-regulatory elements that have the eGene as a target according to the rE2G variant-to-gene prediction model). Only the SV-eQTLs overlapping at least one cCRE are shown; see **Supplementary Table 1** for the complete list.

| SV_ID | CHR | POS | TYPE | LENGTH | AF_ALT | eGENE | P-value | Beta | cCRE(rE2G) |
|---|---|---|---|---|---|---|---|---|---|
| 1-142692086-INS-171 | 1 | 142692086 | INS | 171 | 0.13 | *NBPF15* | 4.73E-09 | -0.35 | 1(0) |
| 1-148521685-DEL-310 | 1 | 148521685 | DEL | 310 | 0.03 | *NOTCH2NLC* | 3.44E-19 | -1.27 | 1(0) |
| 1-148521685-DEL-310 | 1 | 148521685 | DEL | 310 | 0.03 | *SEC22B4P* | 9.42E-21 | -1.24 | 1(0) |
| 2-10041957-DEL-58 | 2 | 10041957 | DEL | 58 | 0.11 | *GRHL1* | 7.61E-09 | -0.35 | 1(1) |
| 2-88741806-INS-3047 | 2 | 88741806 | INS | 3047 | 0.04 | *EIF2AK3* | 4.90E-08 | 0.37 | 1(0) |
| 3-134101117-INS-51 | 3 | 134101117 | INS | 51 | 0.04 | *NUDT16L2P* | 6.58E-09 | 0.46 | 1(0) |
| 4-91252483-COMPLEX-11552 | 4 | 91252483 | COMPLEX | 11552 | 0.02 | *PPM1K* | 3.26E-10 | 0.68 | 15(0) |
| 6-2809526-DEL-70 | 6 | 2809526 | DEL | 70 | 0.2 | *SERPINB6* | 2.83E-08 | -0.29 | 1(1) |
| 7-26232863-DEL-8496 | 7 | 26232863 | DEL | 8496 | 0.19 | *CBX3* | 4.14E-09 | -0.31 | 13(0) |
| 7-136895465-DEL-2185 | 7 | 136895465 | DEL | 2185 | 0.05 | *STMP1* | 1.31E-09 | -0.41 | 2(0) |
| 7-143662575-DEL-21652 | 7 | 143662575 | DEL | 21652 | 0.5 | *TRBV5-1* | 4.70E-11 | 0.35 | 46(0) |
| 7-143662575-DEL-21652 | 7 | 143662575 | DEL | 21652 | 0.5 | *TRBV6-1* | 1.05E-09 | 0.42 | 46(0) |
| 8-1925391-INS-79 | 8 | 1925391 | INS | 79 | 0.17 | *MYOM2* | 1.03E-13 | -0.76 | 1(1) |
| 8-97913536-DEL-65 | 8 | 97913536 | DEL | 65 | 0.67 | *CPQ* | 3.88E-08 | 0.14 | 1(0) |
| 8-145086841-INS-304 | 8 | 145086841 | INS | 304 | 0.29 | *EPPK1* | 5.03E-09 | -0.24 | 1(0) |
| 9-20988220-INS-193 | 9 | 20988220 | INS | 193 | 0.34 | *HACD4* | 1.48E-08 | 0.21 | 1(1) |
| 11-367010-INS-68 | 11 | 367010 | INS | 68 | 0.37 | *NLRP6* | 1.44E-08 | 0.21 | 1(0) |
| 11-367051-DEL-67 | 11 | 367051 | DEL | 67 | 0.33 | *IFITM3* | 7.04E-15 | -0.48 | 1(1) |
| 12-121508103-DEL-65 | 12 | 121508103 | DEL | 65 | 0.04 | *COQ5* | 1.06E-08 | -0.63 | 1(0) |
| 12-131963305-DEL-733 | 12 | 131963305 | DEL | 733 | 0.03 | *EP400P1* | 1.21E-09 | 0.56 | 1(0) |
| 15-87610693-DEL-52 | 15 | 87610693 | DEL | 52 | 0.28 | *ARPIN* | 3.33E-08 | 0.17 | 1(0) |
| 16-94782163-INS-131 | 16 | 94782163 | INS | 131 | 0.46 | *RNF166* | 5.21E-09 | -0.2 | 1(1) |
| 16-95937318-COMPLEX-237 | 16 | 95937318 | COMPLEX | 237 | 0.02 | *VPS9D1* | 4.03E-09 | 0.46 | 1(0) |
| 16-96138787-INS-71 | 16 | 96138787 | INS | 71 | 0.12 | *CICP25* | 1.41E-08 | 0.35 | 1(0) |
| 17-7165549-INS-293 | 17 | 7165549 | INS | 293 | 0.64 | *PHF23* | 2.23E-09 | 0.24 | 1(0) |
| 17-29635045-DEL-677 | 17 | 29635045 | DEL | 677 | 0.09 | *RAB34* | 4.77E-08 | -0.38 | 1(1) |
| 17-48751053-INS-320 | 17 | 48751053 | INS | 320 | 0.04 | *SCRN2* | 3.49E-10 | -0.57 | 1(1) |
| 17-83695580-DEL-86 | 17 | 83695580 | DEL | 86 | 0.26 | *TBCD* | 1.55E-10 | -0.64 | 1(0) |
| 17-84075413-DEL-57 | 17 | 84075413 | DEL | 57 | 0.27 | *B3GNTL1/QTGAL* | 1.46E-08 | -0.2 | 1(1) |
| 18-74731285-DEL-54 | 18 | 74731285 | DEL | 54 | 0.17 | *CNDP2* | 2.08E-17 | -0.59 | 1(0) |
| 19-498725-DEL-159 | 19 | 498725 | DEL | 159 | 0.07 | *GZMM* | 1.65E-09 | -0.34 | 1(1) |
| 19-557247-COMPLEX-302 | 19 | 557247 | COMPLEX | 302 | 0.09 | *HCN2* | 2.60E-08 | 0.64 | 1(1) |
| 19-11311176-DEL-1663 | 19 | 11311176 | DEL | 1663 | 0.08 | *KANK2* | 1.16E-14 | -0.61 | 3(3) |
| 19-54718716-DEL-16422 | 19 | 54718716 | DEL | 16422 | 0.19 | *SIGLEC5* | 4.75E-10 | -0.27 | 13(7) |
| 19-54718716-DEL-16422 | 19 | 54718716 | DEL | 16422 | 0.19 | *SIGLEC14* | 6.44E-12 | -0.49 | 13(7) |
| 19-57188558-INS-77 | 19 | 57188558 | INS | 77 | 0.44 | *OSCAR* | 3.59E-15 | -0.21 | 1(1) |
| 20-64658240-DEL-60 | 20 | 64658240 | DEL | 60 | 0.09 | *TCFL5* | 3.18E-13 | 0.75 | 1(1) |
| 21-40255346-DEL-89 | 21 | 40255346 | DEL | 89 | 0.52 | *PRDM15* | 3.76E-08 | 0.26 | 1(1) |
| 22-25515383-DEL-2605 | 22 | 25515383 | DEL | 2605 | 0.08 | *KIAA1671* | 3.90E-09 | 0.38 | 3(2) |

Next, we used the 1000 Genomes Project (1KGP) SR-WGS dataset genotyped by PanGenie with HPRC2 (16) to replicate the 99 SV-eQTLs identified in CaG. Of these, 93 SV-gene associations (involving 93 SVs and 91 genes) were eligible for replication because the SV had been genotyped by PanGenie in 731 1KGP participants (of which 19% are EUR), and the corresponding gene was present in the 1KGP gene expression matrix. For these eQTL replication analyses, we extracted from the 1KGP dataset and added as covariates the genotypes of variants (SNPs, indels, standard SVs) found in CaG to independently associate with gene expression (**Methods**). We observed a strong replication of the CaG results in the 1KGP data: 66% (61/93) of the associations were in the concordant direction (binomial P-value=0.0035), 41 were nominally significant (P-value <0.05) and 25 remained significant after multiple testing correction (P-value <5.4×10^-4^) (**Supplementary Table 2**).

### SV-eQTLs intersecting ENCODE cCREs

To gain insights into the mechanisms by which the SV-eQTLs might affect gene expression, we assessed the overlap between the independent SV-eQTLs and regulatory elements. We intersected 93 independent SV-eQTLs (six SVs were lost because they have no corresponding coordinates on GRCh38) with ENCODE candidate cis-regulatory elements (cCREs) (26): 36 SV-eQTLs (39%) intersected at least one cCRE, with seven SVs overlapping several due to their large size (**Table 1**). This includes 12 insertions, 21 deletions, and three complex SVs, of which eight deletions and three complex SVs overlapped at least 50% of a cCRE (**Supplementary Figure 7**). SV–eQTLs overlapped regulatory elements more frequently than SVs included in the eQTL analyses (39% vs 23%, odds ratio (95% confidence interval) = 2.13 (1.36-3.28), Fisher’s exact test P-value=0.0007). To further confirm this enrichment, we generated 10,000 sets of 93 randomly selected SVs; for each set, we matched the selected SVs to the SV–eQTLs by SV type and cumulative SV length. Among 10,000 matched random sets, only six sets resulted in ≥36 SV-cCRE overlaps (empirical P-value=0.0006) and only 18 sets resulted in ≥11 SVs overlapping at least 50% of a cCRE (empirical P-value=0.0018) (**Figure 2D-E**). These results indicate that the independent SV-eQTLs identified in CaG using a pangenome-based approach are more likely to overlap regulatory elements than expected by chance.

Next, we used enhancer-gene links data from the ENCODE-rE2G predictive model (28) to assess if the cCRE overlapped by the SV-eQTLs were connected to the same eGenes. We identified 26 cCRE-SV pairs where the regulatory element and the SV were linked and associated, respectively, to the same gene (17 unique SVs and 26 unique cCREs)(**Table 1** and **Supplementary Table 1**). For example, we identified a 1.6 kb deletion in the intron of *KANK2* on chromosome 19 associated with decreased *KANK2* expression (**Figure 3A**). This SV completely deletes two distal enhancers and 35% of another enhancer; these three enhancers are linked to *KANK2* by the ENCODE-rE2G model. We also identified a 677 pb deletion on chromosome 17 associated with decreased *RAB34* expression (**Figure 3B**). The deletion is located 21.3 kb away from *RAB34* (on build GRCh38) and maps to an intron of *SUPTcH*. The deletion removes 74% of a distal enhancer linked to *RAB34* by the ENCODE-rE2G model. Finally, we found a 320 bp insertion on chromosome 17 associated with decreased expression of *SCRN2* (**Figure 3C**). The insertion is in a distal enhancer of *SCRN2* and may affect *SCRN2* expression by disrupting one of its regulatory elements.

**Figure 3.**
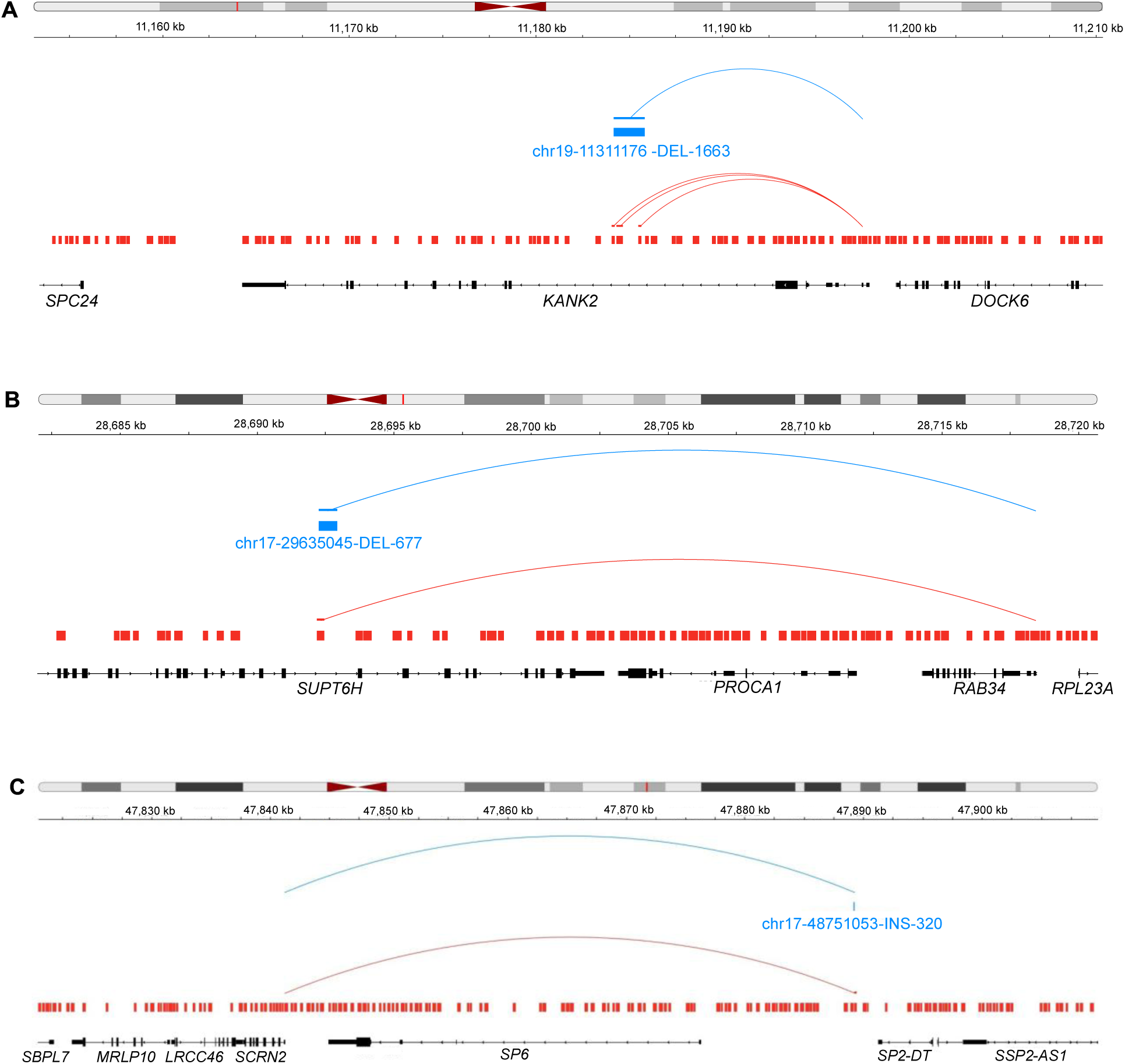
SV-eQTLs overlapping ENCODE cCREs linked to the eGene through the rE2G model predictions. (**A**) A 1.6 kb deletion on chromosome 19 associated with decreased expression of *KANK2* that overlaps three distal enhancers linked to *KANK2*. (**B**) A 677 bp deletion on chromosome 17 associated with decreased expression of *RAB34* that overlaps a distal enhancer linked to *RAB34*. (**C**) A 320 bp insertion on chromosome 17 associated with decreased expression of *SCRN2* that falls in a distal enhancer linked to *SCRN2*. The blue and red boxes correspond to genes and ENCODE cCREs, respectively. The blue lines represent the eQTL association between the SV and the gene (pointing to the gene transcription start site), while the red lines represent the scE2G link between a regulatory element and its target gene. Only the links between the overlapped regulatory elements are shown. The reference genome is GRCh38.

### Multiallelic SV-eQTLs

Pangenomes contain a large proportion of multiallelic variants. In *HPRC2+PanGenie*, 85% of the alternate SV alleles are found at multiallelic loci. Since we used a biallelic representation of the multiallelic variants genotyped by PanGenie, we analyzed in greater depth the SV-eQTLs that implicated an alternate allele of a multiallelic SV. Of the 99 independent SV-eQTLs that we identified, 53 (54%) are multiallelic and include at least two alternate alleles at the same genomic position that were successfully genotyped in CaG by PanGenie. Across the 53 multi-allelic SVs, there were 152 alternate alleles with AF≥1% that were tested in our eQTL analyses. We found that the most common alternate alleles at 36 (68%) of these multi-allelic SVs explained the eQTL signals (**Supplementary Figure 8**). We did not find any cases where multiple alternate alleles of the same SV were associated with gene expression.

Most multiallelic alleles have highly similar sequences, differing only by a few base pairs or by tandem repeat (TR) variations. We focused on a simple form of TR variation in which the alternate alleles contain the same motif but differ in the number of repeats of the motif. At position 104,182,044 on chromosome 10, PanGenie genotyped 49 insertion alleles in CaG, of which 11 were included in the eQTL analyses. The insertions are intergenic and downstream of the gene *INA* (**Figure 4A**). The SV-eQTL allele, associated with a decreased expression of *BORCS7*, has a length of 1,019 bp and an allele frequency of 2.6% in CaG (**Figure 4A**, **Supplementary Table 1** and **Supplementary Table 3**). The most common allele at this locus, a 1,023 bp insertion with a 26% allele frequency in CaG, was not associated with *BORCS7* expression (**Figure 4A**, **Supplementary Tables 3-4**). Using Tandem Repeat Finder (33), a single motif was identified across all 49 alleles, a repetition of the “AC” dinucleotide, with the number of repeats ranging from 15 to 27 (**Supplementary Table 3**). Following multiple sequence alignment, the 11 alleles included in the eQTL analyses were found to differ at five positions: four single-nucleotide differences and differences in the number of motif repeats. The only difference between the most common allele and the SV-eQTL allele is the number of motif repeats. The most common allele contains 20 motif repeats, whereas the SV-eQTL contains 18 (**Supplementary Table 3**). Note also that the SV-eQTL is the allele with the lowest number of repeats among the 11 alleles included in the eQTL analysis (**Supplementary Table 3**). We replicated the association between the SV-eQTL allele and *BORCS7* expression in the 1KGP dataset (**Supplementary Table 5**).

**Figure 4.**
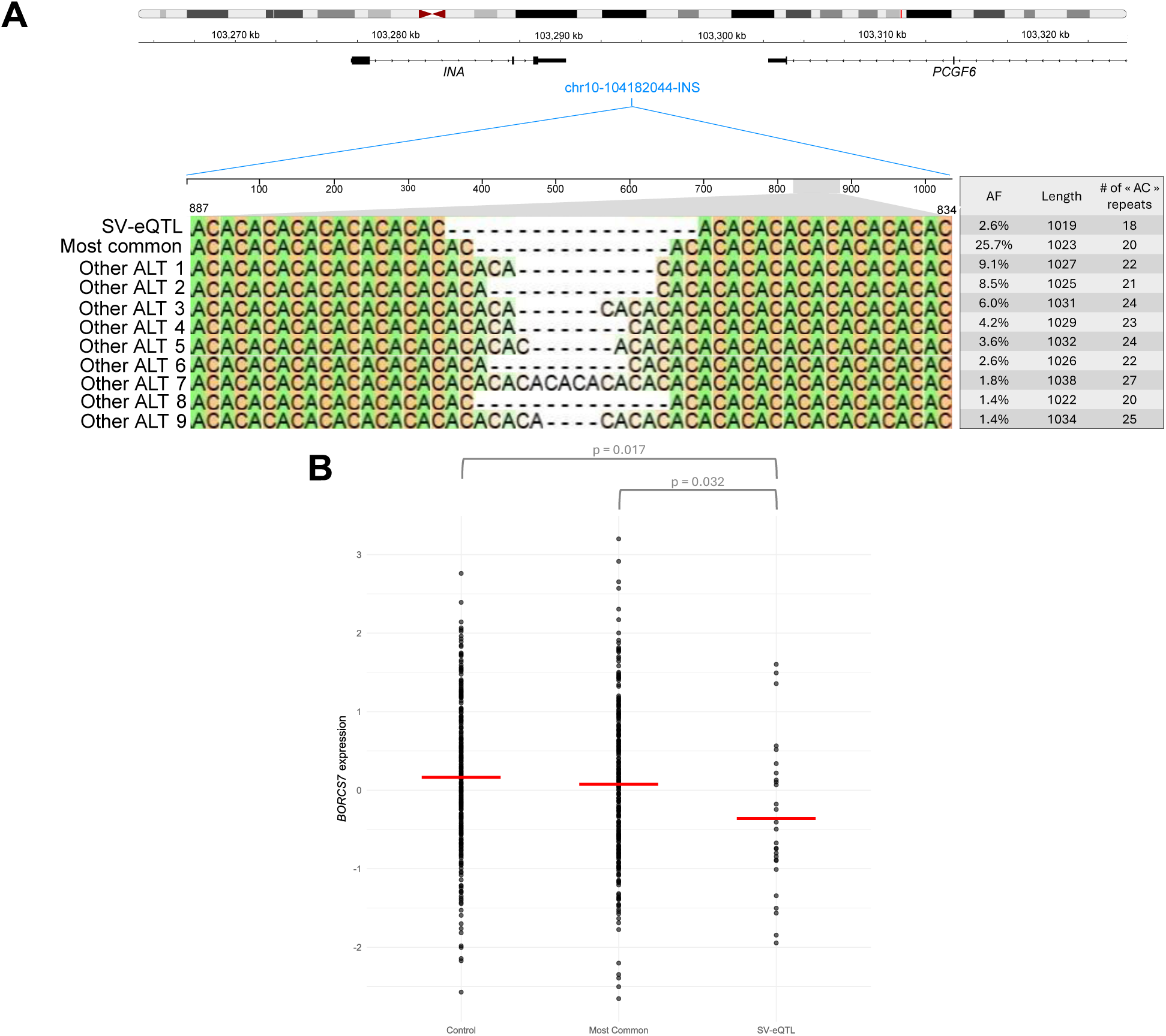
Only one allele at multiallelic locus chr10:104182044 is associated with decreased expression of *BORSC7*. (**A**) Genome view of the multiallelic insertion at chr10:104182044 (top panel). The insertions (blue rectangle) is intergenic, downstream of *INA*, and ∼400kb upstream of *BORSC7*. The reference genome is GRCh38. 49 alternate alleles at this locus were present in the *HPRC2+PanGenie* dataset, of which 11 were successfully genotyped in CaG and included eQTL analyses (AF ≥1%). Multiple sequence alignment of the 11 alleles included in the CaG eQTL analyses (bottom panel). The region of the alignment shown is base pairs 812 to 911 fron the beginning of the inserted sequence, which includes the “AC” repeat notif. The allele associated with the expression of *BORSC7* is naned “SV-eQTL”. (**B**) Expression levels of *BORCS7* in 504 CaG participants, stratified by genotypes at nultiallelic site chr10:104182044. “SV-eQTL” corresponds to the CaG participants carrying one copy of the SV-eQTL allele (N=27). There are no participants carrying two copies of the SV-eQTL allele. “Most connon” corresponds to the CaG participants carrying one or two copies of the nost connon allele (N=238). “Control” corresponds to the renaining participants (N=239). The red lines denote the mean expression level per group. We normalized *BORCS7* expression levels using inverse normal transformation. Two-tail Student’s *t*-test p-values. ALT, alternate allele.

Compared with CaG participants carrying one or two copies of the most common allele (N=238) and those carrying neither the most common allele nor the SV-eQTL allele (control group, N=239), participants carrying one copy of the SV-eQTL allele had higher *BORCS7* expression (**Figure 4B**). *BORCS7* expression did not differ significatively between participants carrying at least one copy of the most common allele and those in the control group (**Figure 4B**). We performed several additional analyses, testing the number of insertion alleles, the cunulative nunber of “AC” dinucleotide repeats across both alleles or their combination but found no significant association with *BORCS7* expression levels (**Supplementary Figure 9** and **Supplementary Table 4**). Together, our results suggest that it is not the insertion allele per se that controls the expression of *BORCS7*, but rather the shorter length (or reduced number of motif repeats) of one alternate allele. Short tandem repeats (STRs) are enriched in cis-regulatory elements in eukaryotes, and changes in STR lengths is often associated with gene expression (34,35). A recent study showed that STRs length dictates local transcription factor binding affinity and recruitment capacity, thereby modulating gene expression (35). However, further investigation is needed to confirm whether this transcription factor tuning mechanism drives the association between the shortest insertion allele and *BORCS7* expression.

### Association of SV-eQTLs with human quantitative traits

We investigated whether the independent *HPRC2+PanGenie* SV-eQTLs were associated with clinically relevant human quantitative phenotypes. In CaG, we tested the 99 SVs against 42 quantitative traits (e.g. hematological and anthropometric traits, blood lipid levels) but found no associations that reached the statistical significance threshold after multiple testing correction (P-value<1.2×10^-5^) (**Supplementary Table 6**). Given the limited statistical power of our cohort (N=2,173), we decided to assess whether the SV-eQTLs were in LD with small variants associated with complex human phenotypes by GWAS. In the CaG dataset, we identified SNP and indel proxies genotyped by PanGenie from HPRC2 that were in strong linkage disequilibrium (r^2^ ≥0.8) with the 99 SV-eQTLs. We found 300 unique small variant proxies for 65 SVs. We queried these small variants against the Open Targets fine-mapping credible sets database (27) and found 94 unique proxies corresponding to 36 SVs that are included in 2,566 different fine-mapped credible sets. These GWAS credible sets are implicated in phenotypes such as Parkinson’s disease, fatty acid levels, neuroticism, albumin levels and blood pressure (**Supplementary Table 7**). For instance, the SV-eQTL associated with *KANK2* expression (**Figure 3A**) is in perfect LD (r^2^=1) with fine-mapped GWAS variants associated with red blood cell traits, triglyceride levels and blood pressure (**Supplementary Table 7**). These results suggest that SV-eQTLs in strong LD with small variants from the Open Targets credible sets represent candidate causal variants for these complex human phenotypes.

## DISCUSSION

LR-WGS technologies together with graph genome methodologies are revolutionizing how we study the impact of SVs on human phenotypic variation. While high-quality telomere-to-telomere (T2T) personal genome assemblies remain the gold-standard to fully capture all types of genetic variation, methods have been developed that leverage pangenomes and SR-WGS data to gain new insights into SVs. By combining the software PanGenie together with phased haplotypes from HPRC, we increased by 25-fold the number of SVs that we could genotype in CaG participants when compared to SV detection that relies solely on SR-WGS data and a linear reference genome. Moreover, this pangenomic approach makes it possible to genotype SVs that are not in sufficient linkage disequilibrium with small variants to be included in SV imputation panels. We showed that these pangenome-derived SVs independently associate with gene expression, are enriched for regulatory elements and are in LD with fine-mapped GWAS signals. This result highlights the importance to consider SVs in planning post-GWAS functional experiments and motivate the development of newer genome editing methods to precisely engineer SV breakpoints and rearrangements in cellular or animal models.

More than half of the SV-eQTL that we identified by conditional analyses were multiallelic variants. In contrast to our expectations, we found many such multiallelic SVs where alleles of intermediate frequency, but not the most common alleles, were associated with gene expression (**Supplementary Figure 8**). Furthermore, we did not identify multiple alleles of the same multiallelic SVs that were associated with gene expression. Based on these observations, we suspect that our strategy to biallelize multiallelic variants prior to association testing is suboptimal. Testing each allele one-by-one increases the number of tests and decreases power, but can be sufficient when only one of the alleles is functional (as in the example presented in **Figure 4**) or if only one of the causal alleles is common. However, if >1 alleles are functional and common, the biallelization method lowers power because it includes the alternate causal allele(s) in the reference genotype. Tests that collapse multiple variants together, such as those designed for rare variant testing (i.e. burden, kernel and omnibus methods) might provide an alternative approach that deserve further methodological development (36). With the use of pangenome-based approaches gaining in popularity and the prevalence of multiallelic variants in those resources, we need to develop better statistical models that can account for multiallelic variants in association testing.

Our project has several limitations. First, because we used a genotyping approach, we could only detect SVs represented in HPRC1-2. Consequently, SVs specific or enriched in CaG were missed. Second, we did not have access to a well-matched cohort to replicate the whole-blood eQTLs found in the mostly European-ancestry CaG cohort. Indeed, the 1KGP dataset used in our replication effort included 81% of non-European-ancestry participants and gene expression was measured in lymphoblastoid cell lines. Accordingly, there may be true positive SV-eQTLs found in CaG that did not replicate in 1KGP due to these differences. Third, because there are currently no large genetic datasets with genotypes for both small variants and SVs, we could not perform colocalization analyses to confirm the concordance of the GWAS (Open Targets) and eQTL signals. Such analyses will await the genotyping of SVs in large datasets, such as the UK Biobank or All of Us Research Program, that are well-powered for GWAS.

## CONCLUSIONS

Our study provides but a glimpse of the impact that SVs might play in shaping the architecture of human phenotypes. Indeed, since the genetic ancestry of the CaG participants differ from the reference populations currently included in HPRC, it is likely that the number of SVs that we genotyped is low in comparison with the number of SVs that we could recover if a Quebec-specific pangenome was available. Thus, although global pangenome efforts are extremely valuable, comprehensive study of genetic variation, including SVs, will require the creation of population-specific pangenomes in parallel.

## METHODS

### CaG short-read whole-genome sequencing data

The CaG SR-WGS dataset and SV calling using standard SR-WBS-based methods has been described elsewhere (24). 2,184 unrelated participants were selected for sequencing. DNA samples were sequenced on Illumina NovaSeq sequencers using PCR-free libraries and a paired-end protocol. The Illumina DRAGEN pipeline was used for variant calling. After quality control, 2,173 samples and 80,407,530 biallelic variants were left. We called SVs in each participant using DELLY and Manta and merged the two SV sets using SURVIVOR (37–39). We genotyped 25,820 SVs (24,022 deletions and 1,798 duplications) and generated a multi-sample VCF using muCNV (40). We used GRCh38 as reference genome for these analyses.

### Genotyping variants from the HPRC pangenomes

We used PanGenie v4.1.1 (29) to genotype variations from HPRC2 into 2,173 CaG participants with SR-WGS data. We used the PanGenie-ready VCFs from the Minigraph-Cactus graph with CHM13 as the reference sequence. We first ran the preprocessing step with the command PanGenie-index. We then genotyped each sample separately. To optimize computational resources, we ran the genotyping command in two steps. The first step (command PanGenie with –w option) was run using 24 threads (options -j 24 and -k 24). The second step, where the output VCF is written, uses only one core (command PanGenie-vcf). We used the default number of haplotypes, 15, for the haplotype-sampling step of the pipeline, as recommended. We used bcftools merge (41) to combine the samples’ VCFs output by PanGenie into a multi-sample VCF. PanGenie output a total of 40,255,343 variants from HPRC2. PanGenie computes genotypes for all bubbles in the graph. To convert the bubble genotypes into variant genotypes and extract all the nested variant alleles represented in the bubbles, we used the script convert-to-biallelic.py provided on the PanGenie GitHub page. This converted the multiallelic VCF into a biallelic VCF representing variant alleles. We filtered out monomorphic variants (with AF=0 or AF=1) using bcftools filter, which left us with 34,226,043 biallelic variants. We added a new INFO tag corresponding to the lengths of each allele (LEN_REF and LEN_ALT) using bcftools annotate. We created unique IDs for each variant with a home-made Python script. To extract the SVs, we filtered the VCF based on the variant’s allele length using bcftools filter. We defined an SV as a variant that has at least one allele (REF or ALT) with a length ≥50 bp (9). We further defined three types of SVs: deletions (LEN_REF ≥ 50 and LEN_ALT = 1), insertions (LEN_REF = 1 and LEN_ALT ≥ 50) and complex SVs ([LEN_REF > 1 and LEN_ALT ≥ 50] or [LEN_REF ≥ 50 and LEN_ALT > 1]). We extracted 645,198 SVs, including 103,630 deletions, 455,071 insertions and 86,979 complex SVs.

We used PanGenie v3 to genotype HPRC1 variants into the 2,173 CaG participants with SR-WGS data. We used the PanGenie-ready VCFs from the Minigraph-Cactus graph with CHM13 as the reference sequence. We executed the same steps as mentioned above but we did not split the genotyping command in two steps since PanGenie v3 has no implementation of the haplotype-sampling step. PanGenie genotyped 27,829,007 biallelic variants, including 318,162 SVs (68,161 deletions, 171,703 insertions, and 77,298 other complex SVs).

### Number of SVs per genome and per ancestral group

We used PLINK2 (42,43) --sample-counts function to calculate the number of observed variants per sample. We excluded variants in the non-PAR regions of chromosome X. We defined the number of SVs per genome as the sum of the number of homozygous alternate genotypes and the number of heterozygous genotypes.

### Concordance of PanGenie genotypes

We used LR-WGS data of 20 CaG participants generated as part of another project using Pacific Biosciences HiFi technology (∼30X coverage) (30). The 20 samples are of French-Canadian ancestry (QFC), have their four grandparents born in Canada, and have French as their first spoken language. A graph was made combining the 40 assembled CaG haplotypes and the 464 HPRC2 haplotypes (HPRC2-CaG graph). It was then projected into a VCF format using Minigraph-Cactus (44). To make it comparable to the VCF generated by PanGenie in terms of bubble decomposition, we processed the HPRC2-CaG graph VCF with the prepare-vcf-MC-pipeline (available on the PanGenie GitHub page), which is the pipeline used to make the VCFs from graphs compatible with the VCF input that PanGenie requires. The Y chromosome was removed from the HPRC2-CaG graph VCF before executing the pipeline. The pipeline produced two VCFs: a multiallelic VCF representing bubbles and a biallelic VCF. We used the biallelic VCF for subsequent analyses. We subset both the HPRC2–CaG graph VCF and the *HPRC2+PanGenie* VCF to retain the 20 CaG samples with available LR-WGS data. We then filtered out monomorphic variants and extracted SVs as mentioned above. The number of SVs from each dataset was comparable, with 40,995 deletions, 87,229 insertions and 13,617 complex SVs from the *HPRC2-CaG-LR* dataset, and 44,050 deletions, 88,341 insertions and 13,947 complex SVs from the *HPRC2+PanGenie* dataset. To measure genotype concordance, we focused on SVs that were identical between the two datasets: 11,081 deletions, 17,038 insertions and 476 complex SVs, for a total of 28,595 SVs. We intersected the two datasets, processing each type of SV separately, using bedtools intersect v2.31.0 (options -F 1 -f 1) (45). For insertions, we kept the intersecting pairs that had the same inserted sequence. For complex alleles, we did not use bedtools, but we created identical pairs by matching their position, the REF allele, and the ALT alleles. We created plink files for each dataset, containing only the SVs that were paired (with option -- vcf-half-call “missing” for the *HPRC2-CaG-LR* dataset). We used the PLINK --pgen-diff function to calculate the number of discordant genotypes between the paired SVs (43,46). We defined the genotype concordance rate as the number of concordant genotypes divided by the total number of compared genotypes.

We also calculated genotype concordance for the *HPRC1*+*PanGenie* dataset and the S*tandard SV* dataset. We identified 36,166 deletions, 51,257 insertions and 21,062 complex SV from the *HPRC1*+*PanGenie* dataset that were genotyped in the 20 CaG samples with available LR-WGS data. We matched 2,081 deletions, 2,202 insertions and 210 complex SVs between the *HPRC1*+*PanGenie* dataset and the *HPRC2-CaG-LR* dataset, and we used these matches to compute genotype concordance as mentioned above. For the S*tandard SV* dataset, we identified 6,494 deletions in the 20 CaG samples with available LR-WGS data, and we matched 850 of them with deletions from the *HPRC2-CaG-LR* dataset.

### eQTL analysis

RNAseq data from whole blood was generated for 911 CaG participants, of which 504 have SR-WGS data and have passed quality-control, including 471 QFC, 5 QHA and 3 QMO samples (24,47). Quality control and processing of CaG RNAseq data have already been described (24). The PEER factors were calculated from the normalized LCPM gene counts, and the gene expressions were inverse normal transformed.

We used tensorQTL v1.0.10 (32) to test the association between SVs and gene expression We used both the *HPRC2+PanGenie* dataset and the S*tandard* SV dataset. We ran tensorQTL separately for the two datasets, since the S*tandard* approach SVs are on build GRCh38 and the HPRC2 SVs are on build CHM13. We tested SVs that passed quality control filters of genotype nissingness ≤10% and MAF ≥1% in the 504 CaG participants. This left 161,624 SVs from the *HPRC2+PanGenie* dataset and 4,653 SVs from the S*tandard* approach. We used age, sex, the first 10 PCs computed from the genetic data and 60 PEER factors as covariates. TensorQTL requires a bed file with the transcription start sites (TSS) of each gene. We extracted the TSS of the genes for which expression data was available from the CHM13 or GRCh38 genome annotation files. By default, tensorQTL uses a 1 Mb window around the given TSS (TSS ± 1Mb, cis windows). TensorQTL does not analyze phenotypes for which there are no variants in the specified window, which left us with 13,558 genes in CHM13 and 12,991 genes in GRCh38. We did not test chromosome Y nor the mitochondrial genome.

To evaluate the added value of a pangenomic approach before association analysis, we tested the association between SVs from the *HPRC2+PanGenie* dataset and gene expression while conditioning on variants identified from the CaG SR-WGS data using the *Standard* approach (i.e. based on GRCh38). We first conducted linear regressions using PLINK2 (42,43) between gene expression and S*tandard* variants (SNPs, INDELs and SVs) from the 504 CaG participants with SR-WGS and RNAseq data. We used age, sex, the first 10 PCs computed from the genetic data and 60 PEER factors as covariates. We then gave the results of the linear regressions to a home-made Python script. This script conducts multiple rounds of conditional analyses. It first identifies the *Standard* approach variant from the linear regression with the smallest P-value and conducts a conditional analysis using this variant as an additional covariate. It repeats the process with the top-most associated variant until no variant remains genome-wide significant. From this script, we extracted all the variants that were conditioned on. We used PLINK2 to perform linear regressions between gene expression and SVs from *HPRC2-PanGenie* and added the participants’ genotypes to the list of conditioned variants identified by the Python script to the basic covariates used in the models.

### Intersection of SV-eQTLs with ENCODE candidate cis-regulatory elements

We used the ENCODE Registry of candidate cis-Regulatory Elements (cCREs) (Registry V3) (26). Since there were no ENCODE cCREs files with coordinates on CHM13, we lifted over the coordinates of the SVs tested in the eQTL analyses using the UCSC LiftOver online interface with the default parameters (48). 137,624/161,654 SVs had coordinates on GRCh38, including 93/99 SVs-eQTLs. We intersected the cCREs and the SVs using bedtools intersect v2.31.0 (45). For complex SVs, we used the coordinates of the reference allele. To confirm the enrichment of SV–eQTLs overlapping cCREs compared with all tested SVs, we created 10,000 random sets of SVs. For each set, we randomly selected SVs matched to the SV–eQTL set based on the SV type and cumulative length. Among the SV–eQTLs with coordinates available in GRCh38, 48 were insertions, 39 were deletions, and 6 were complex SVs, with the deletions and complex SVs spanning a cumulative length of 73,454 bp. Therefore, for each set, we randomly selected 48 insertions and 45 deletions and/or complex SVs, ensuring that their cumulative length differed by no more than 10% from that of the SV– eQTLs. We then intersected the selected SVs with cCRE annotations and counted the number of unique SVs overlapping at least one cCRE and at least 50% of a cCRE. P-values were calculated as the proportion of permutations with an overlap equal to or greater than the observed number of SV–eQTLs: (1) the number of sets with ≥36 SVs overlapping at least one cCRE divided by 10,000, and (2) the number of sets with ≥11 SVs overlapping at least 50% of a cCRE divided by 10,000. We used the cCRE-Gene-Links data generated by the ENCODE-rE2G model to link the intersected elements to a target gene (28).

### Annotation of SV-eQTLs with snpEff

We used snpEff v5.4 to annotate the SV-eQTLs (49). We first created the database for build CHM13 (command build, with options -noCheckCds and -noCheckProtein). We then used this database to annotate the VCF.

### Multiallelic SV-eQTLs

We used Tandem Repeat Finder version v4.09 with the recommended parameters (2 5 7 80 10 50 2000) to identify the SV-eQTLs that were tandem repeats variations (33). For the multiple sequence alignment of the alternate alleles at locus chr10:104182044, we used the EMBL-EBI Clustal Omega web tool (50–52).

### Association of SVs with quantitative traits in CaG

We used PLINK2 (42,43) to perform linear regressions (--glm function) and test the association between the 99 independent SV-eQTLs and 42 quantitative phenotypes. The phenotypes were rank-inverse normal transformed on the entire CaG cohort, as previously described (24). We used age, sex and the first 10 PCs as covariates. The covariates were normalized with the option --covar-variance-standardized. Since we tested 99 SVs and 42 phenotypes, we defined the significance threshold as P-value <1.2×10^-5^ (0.05 / (99 x 42)).

### Small variant LD proxies tagging SV-eQTLs

We extracted small variants (SNPs and indels) from the *HPRC2+PanGenie* dataset using bcftools filter (41) by filtering on the lengths of the alleles (LEN_REF <50 and LEN_ALT <50) and then generated plink files. In parallel, we generated a set of plink files containing only the 99 independent SV-eQTLs. After merging the plink file sets, we calculated LD between SV-eQTLs and small variants with a MAF ≥1% using the PLINK function --r2 (43,46). We defined a small variant proxy as a small variant with a LD r^2^ ≥0.8 with an SV-eQTL. We lifted over the small variant proxies from CHM13 to GRCh38 using the UCSC liftover tool (48). We queried the small variants in the Open Targets credible set database using the Python API PySpark 4.1.1 (27). To obtain the phenotypes and the predicted genes associated with the credible sets, we used the GWAS/QTL Study database and the Locus-to-Gene (L2G) prediction database.

### SV-eQTLs replication in the 1000 Genomes Project dataset

We used the 1000 Genomes Project (PanGenie+HPRC2+1KGP) dataset (16) to replicate the associations between gene expression and the 99 independent SVs identified through our conditional eQTL analyses in CaG. We first extracted the SVs from the PanGenie+HPRC2+1KGP dataset based on the lengths of the REF and ALT alleles using bcftools and queried the SV-eQTLs, matching them exactly by positions and alleles. We identified 97/99 SV-eQTLs in the PanGenie + HPRC2 + 1KGP dataset. We used the 1KGP gene expression data from the Multi-Ancestry Analysis of Gene Expression (MAGE), which comprises RNAseq data from lymphoblastoid cell lines derived from 731 individuals (N_AFR_=196, N_AMR_=113, N_EAS_=141, N_EUR_=142, N_SAS_=139) (53). We found 91/101 eGenes in the 1KGP gene expression data, which left us with 93 SV-Gene pairs to try to replicate. Since the SV-eQTLs were identified through conditional analysis on variants detected using the *standard* approach, for each gene, we listed the small variants from CaG that were conditioned on and queried them in the 1KGP GRCh38 dataset (54–56). For 6/91 genes, we did not find any of the conditioned variants in the 1KGP dataset. Note that we also did not consider two SVs that were listed as the conditioned variants since they are not present in the 1KGP small variants datasets. We then tested the association between each SV-Gene pairs in the 731 1KGP participants using PLINK2. We used the covariates from MAGE, which includes sex, the 5 first genetic PCs and 60 PEER factors, and added, for each gene, the genotypes of the successfully queried small variants. The significance threshold corrected for the number of tests is P-value=0.05/93=5.4×10^-4^.

### Public Data Accession

The PanGenie-ready VCFs from the Minigraph-Cactus graph with CHM13 as the reference sequence for HPRC1 and HPRC2 were respectively obtained from (57) and (58). The prepare-vcf-MC-pipeline used on the HPRC-CaG graph VCF is available from (59). The ENCODE cis-regulatory elements and the cCRE-Gene-Links data generated by the ENCODE-rE2G mode were downloaded from the SCREEN (Search Candidate cis-Regulatory Elements by ENCODE) portal (60). The sequence, protein and genes files for the annotation with snpEff were downloaded from (61). The Open Targets databases we used are available for download (62). The vcf for the PanGenie + HPRC2 + 1KGP dataset was obtained from (63). The 1KGP gene expression data from the MAGE project (53) and the 1KGP covariates used for replication were obtained from(64). The 1KGP GRCh38 dataset (54–56) was obtained from (65).

## DECLARATIONS

## Ethics approval

The CARTaGENE cohort (CaG) is approved by the Ethics Committee of the CHU Ste-Justine and participants were recruited in accordance with the Declaration of Helsinki. This project is approved by the Ethics Committee of the Montreal Heart Institute (Project #2017-2247).

## Consent for publication

Not applicable.

## Availability of data and materials

Access to the CARTaGENE (CaG) genetic and phenotypic data is controlled in accordance with the consent provided by participants. The procedure for requesting access to CaG data is described at https://portail.cartagene.qc.ca/data-access-process. Access requests must be submitted through the CaG online system and must include the following documents: a completed application form (online), a research protocol, a REB approval letter, and a CV. Applications are reviewed by the CaG Data Access Committee (DAC) to ensure compliance with participant consent and CaG access policies. Applicants are generally informed of the decision within approximately 1–2 months, depending on the completeness of the application and the review process. Questions regarding data access can be addressed to. If access is granted, data may be used only within the scope of the approved research project and in accordance with CaG access conditions.

The code used for data analysis and the generation of figures has been deposited on GitHub (66) and is available on Zenodo (67).

## Author’s contributions

Conceived and designed the analyses: R.L. and G.L.; Collected the data: R.L. and C.G.; Performed analyses: R.L.; Secured funding and supervised the work: G.B. and G.L.; Wrote the manuscript: R.L. and G.L., with contributions from all authors.

## Acknowledgements

We thank all CARTaGENE participants who contributed data to this study. We also thank Jean-Christophe Grenier and Julie Hussin for processing the RNA-seq data.

