## Supplementary Figures for "A pangenomic approach identifies new structural variants associated with gene expression in the CARTaGENE cohort"

**Supplementary Figure 1. Number of SVs genotyped by each approach in 2,173 CaG participants.** The SVs were genotyped using PanGenie and HPRC1 or HPRC2 in CaG participants with short-read whole-genome sequencing data. SVs were also detected in the same participants using a *Standard SV* detection approach based on the GRCh38 reference genome (Standard). The x-axis represents the approach used to genotype SVs. The y-axis represents the SV counts. PanGenie does not distinguish between insertions and duplications. There are no insertions in the *Standard SV* dataset. COMPLEX, complex SVs. DEL, deletions. INS/DUP, insertions and/or duplications.

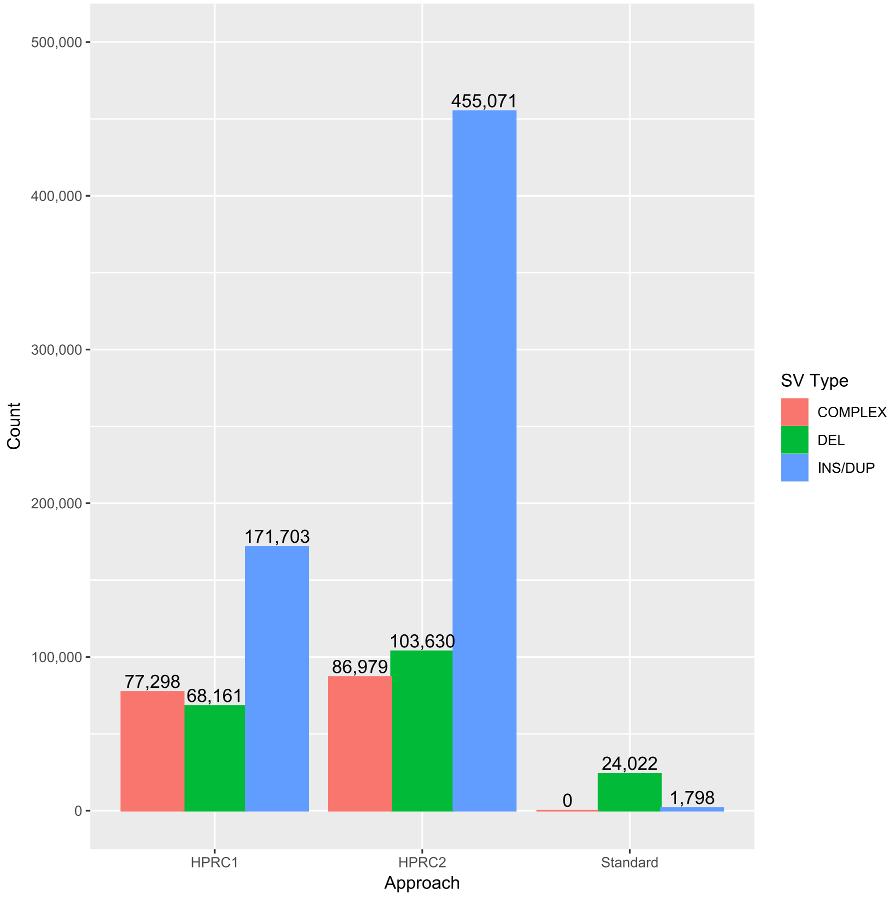

**Supplementary Figure 2. Distribution of SV allele frequencies in 2,173 CaG participants.** The allele frequencies of SVs from the (**A**) *HPRC1+PanGenie* dataset, (**B**) the *HPRC2+PanGenie* dataset and (**C**) the *Standard SV* dataset were calculated in three ancestral groups (x-axis): CARTaGENE participants of French-Canadian ancestry (QFC, N=1,756), CARTaGENE participants of Haitian ancestry (QHA, N=163), and CARTaGENE participants of Moroccan ancestry (QMO, N=131). "ALL" corresponds to the allele frequencies calculated in the 2,173 CARTaGENE participants. The colors represent the allele frequency bin, while the y-axis represents the proportion of SVs in each bin.

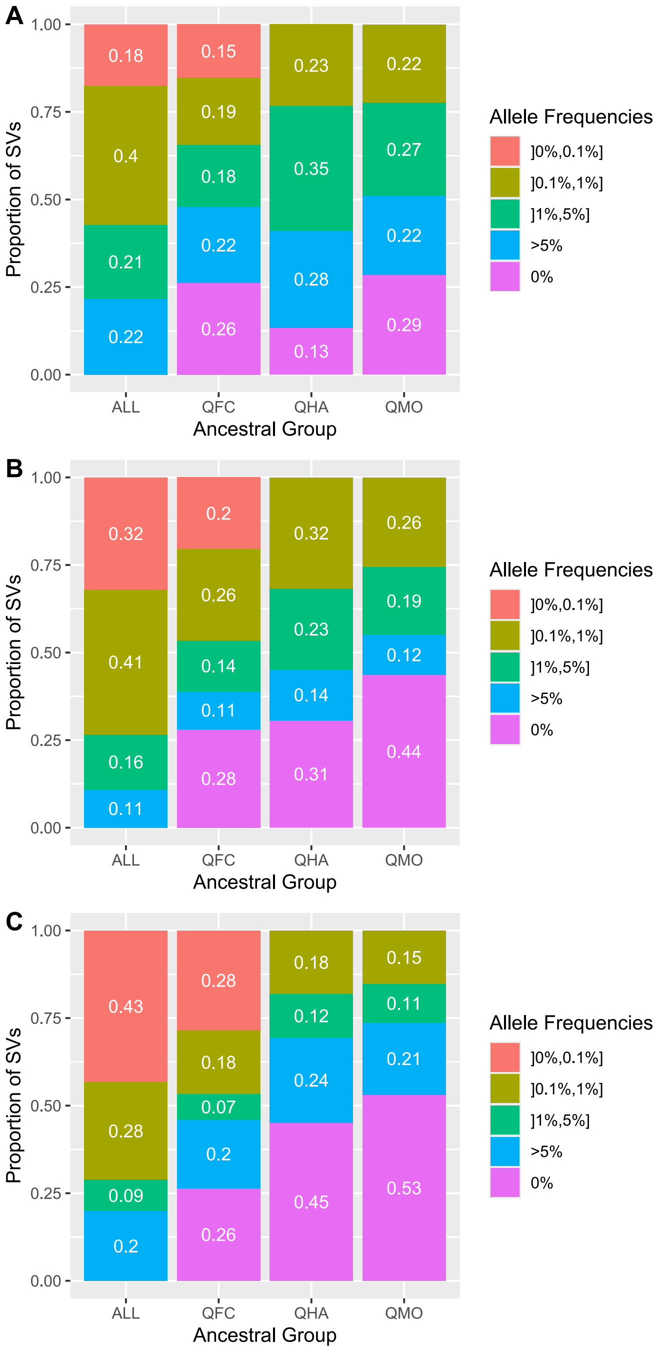
A
B
C
Proportion of SVs
1.00
0.75
0.50
0.25
0.00
ALL
QFC
QHA
QMO
Ancestral Group
Allele Frequencies
]0%,0.1%]
]0.1%,1%]
]1%,5%]
>5%
0%
0.18
0.4
0.21
0.22
0.15
0.19
0.18
0.22
0.26
0.23
0.35
0.28
0.13
0.22
0.27
0.22
0.29
0.32
0.41
0.16
0.11
0.2
0.26
0.14
0.11
0.28
0.32
0.23
0.14
0.31
0.26
0.19
0.12
0.44
0.43
0.28
0.09
0.2
0.28
0.18
0.07
0.2
0.26
0.18
0.12
0.24
0.45
0.15
0.11
0.21
0.53

**Supplementary Figure 3. Association between gene expression and SVs in 504 CaG participants.** Manhattan plot of the association (**A**) between the expression of 13,558 genes and 161,624 SVs from the *HPRC2+PanGen*ie dataset and (**B**) between the expression of 12,991 genes and 4,653 SVs from the *Standard SV* dataset. The red dashed lines represent the threshold used to denote genome-wide significance for cis-eQTL (P = $5 \times 10^{-8}$).

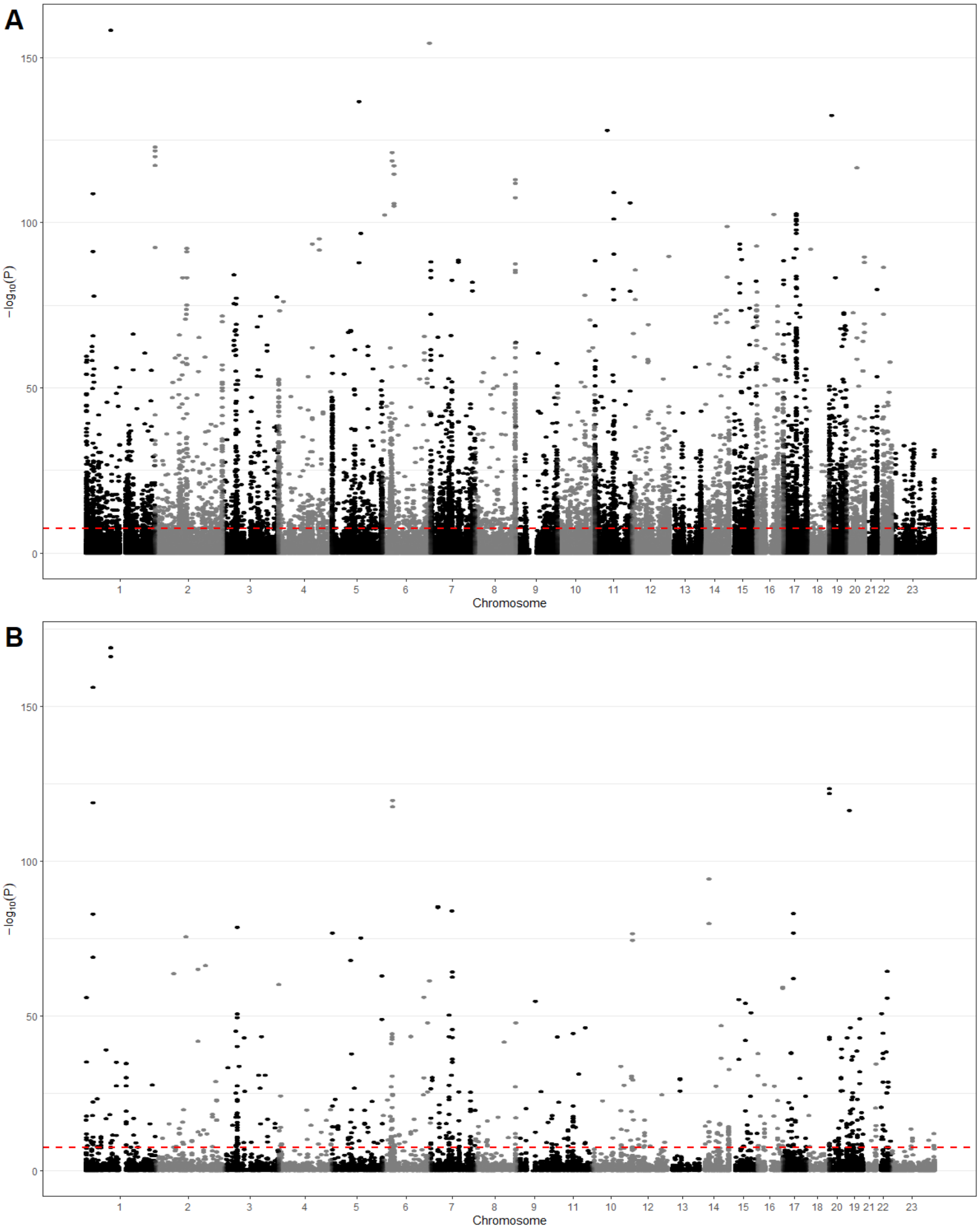

**Supplementary Figure 4. P-value comparison of the independent *HPRC2+PanGenie* SV-eQTLs before and after conditional analyses on variants detected using the *Standard* approach.** The P-values of the 99 independent SV-eQTLs (x-axis) before and (y-axis) after the conditional analysis on variants detected using the Standard approach (SR-WGS, GRCh38). The colors represent the significance of the SVs before the conditional analysis. The red dotted lines represent the threshold used to denote genome-wide significance for cis-eQTL (P-value=$5x10^{-8}$).

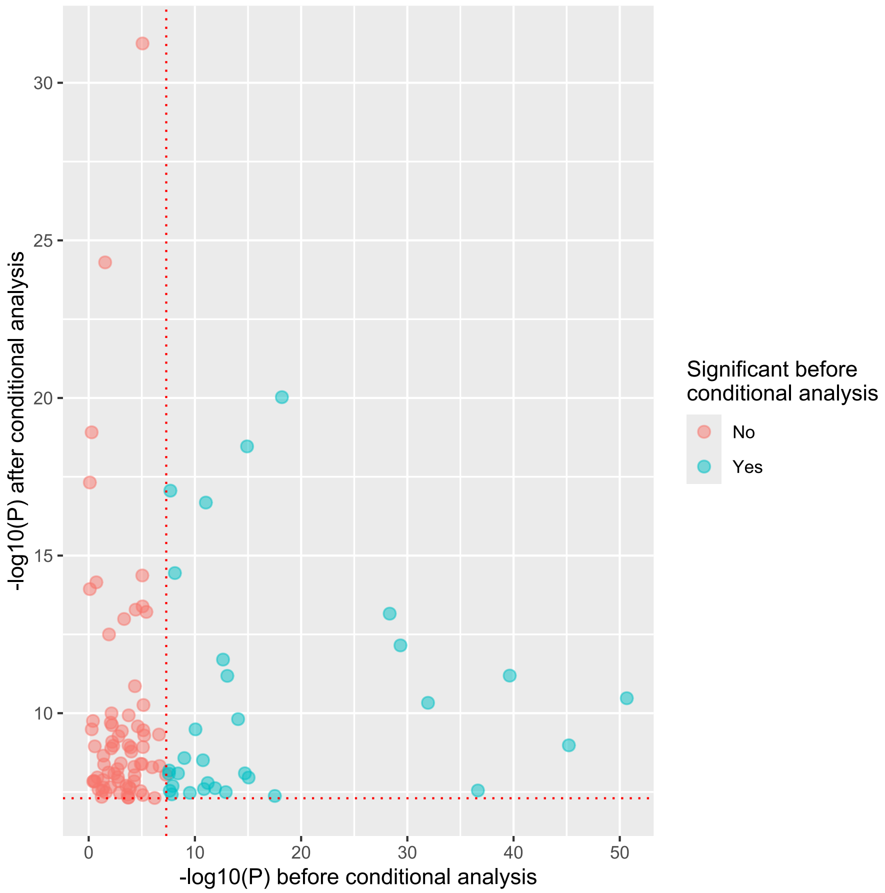

**Supplementary Figure 5. Comparison of allele frequencies between *HPRC2-PanGenie* SVs tested in the eQTL analysis and SVs independently associated with gene expression (cis-eQTLs).** 161,624 SVs from the *HPRC2+PanGenie* dataset were tested in the eQTL analyses, and 99 independent SV-eQTLs were identified after conditional analyses. The alternate allele frequencies (x-axis) were calculated in 2,173 CaG participants. The y-axis displays the density relative to the group (SV-eQTLs (red) or all tested SVs (blue)). Wilcoxon rank-sum test P-value=$2.1\times10^{-15}$. ALT, alternate.

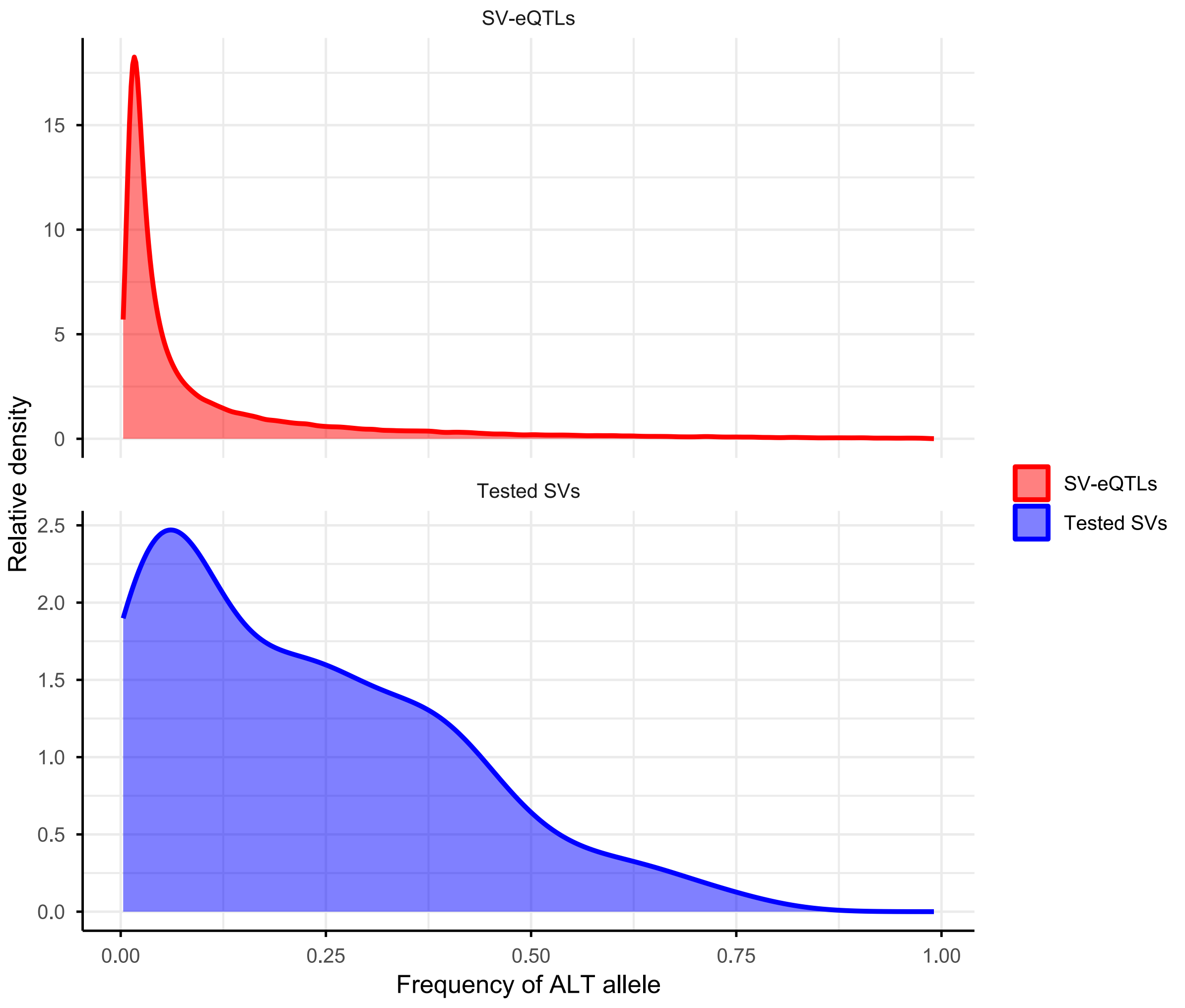

**Supplementary Figure 6. Comparison of genotype quality (GQ) between SVs included in the eQTL analyses and SV-eQTLs.** 161,624 SVs from the *HPRC2+PanGenie* dataset were included in the eQTL analyses, and 99 independent SV-eQTLs were identified after conditional analyses. (**A**) The GQ generated by PanGenie of all SV genotypes. The colors represent the SV sets. The x-axis represents the GQ and the y-axis its proportion in the SV set. (**B**) Comparison of the proportion of SV genotypes with GQ ≥ 30 (green) and GQ < 30 (red) for the 99 SV-eQTLs (left) and the 161,624 tested SVs (right). Pearson's Chi-square P-value $<2\times10^{-16}$.

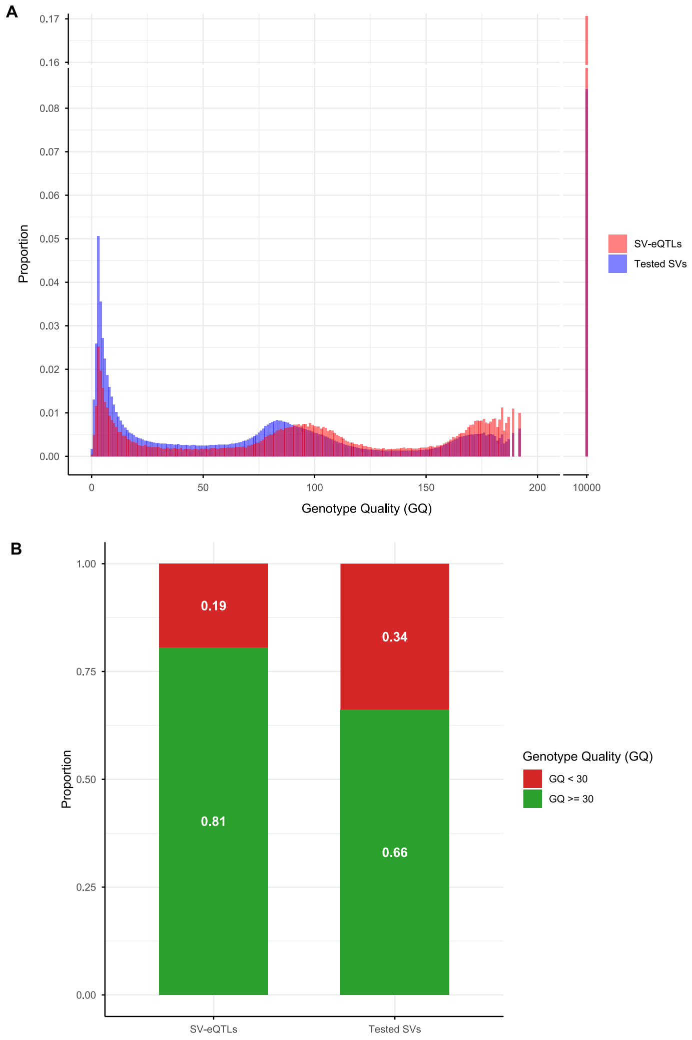

**Supplementary Figure 7. ENCODE candidate cis-regulatory elements (cCREs) overlapped by the SV-eQTLs.** Each bar represents a cCRE overlapped by a deletion or a complex SV. The x-axis represents the proportion of the cCRE that is covered by the SV. The color of the bars represents the cCRE class. The last number in the SV identifier corresponds to its length in base pairs (length of the reference allele for complex SVs). The positions and the lengths in the identifier correspond to those on build CHM13. CA, chromatin accessibility. TF, transcription factor.

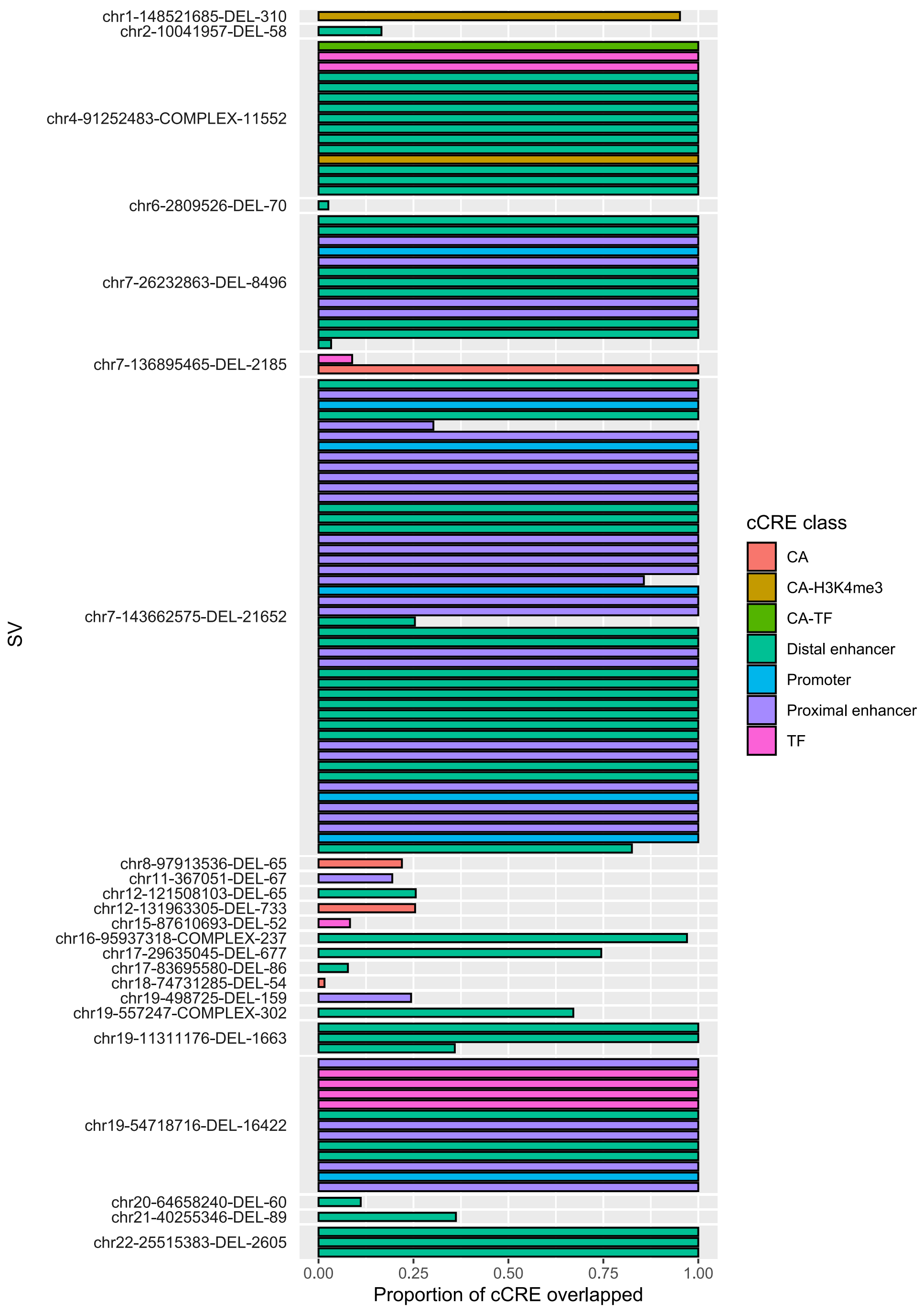

chr1-148521685-DEL-310
chr2-10041957-DEL-58
chr4-91252483-COMPLEX-11552
chr6-2809526-DEL-70
chr7-26232863-DEL-8496
chr7-136895465-DEL-2185
chr7-143662575-DEL-21652
chr8-97913536-DEL-65
chr11-367051-DEL-67
chr12-121508103-DEL-65
chr12-131963305-DEL-733
chr15-87610693-DEL-52
chr16-95937318-COMPLEX-237
chr17-29635045-DEL-677
chr17-83695580-DEL-86
chr18-74731285-DEL-54
chr19-498725-DEL-159
chr19-557247-COMPLEX-302
chr19-11311176-DEL-1663
chr19-54718716-DEL-16422
chr20-64658240-DEL-60
chr21-40255346-DEL-89
chr22-25515383-DEL-2605
SV
0.00
0.25
0.50
0.75
1.00
Proportion of cCRE overlapped
cCRE class
CA
CA-H3K4me3
CA-TF
Distal enhancer
Promoter
Proximal enhancer
TF

**Supplementary Figure 8. Frequency of the alternate alleles of the multiallelic SV-eQTLs.** The y-axis represents the loci of the associated SV-eQTLs that are multiallelic, that is SV with other alleles at the same genomic position in the *HPRC2+PanGenie* dataset. The x-axis represents the allele frequency of the alleles in 2,173 CaG participants. The blue dots represent the SV-eQTL-associated alleles, while the black and grey dots represent, respectively, the tested and untested other alternate alleles at the same loci. The red line denotes the 1% minor allele frequency threshold for SV testing. AF, allele frequency. ALT, alternate.

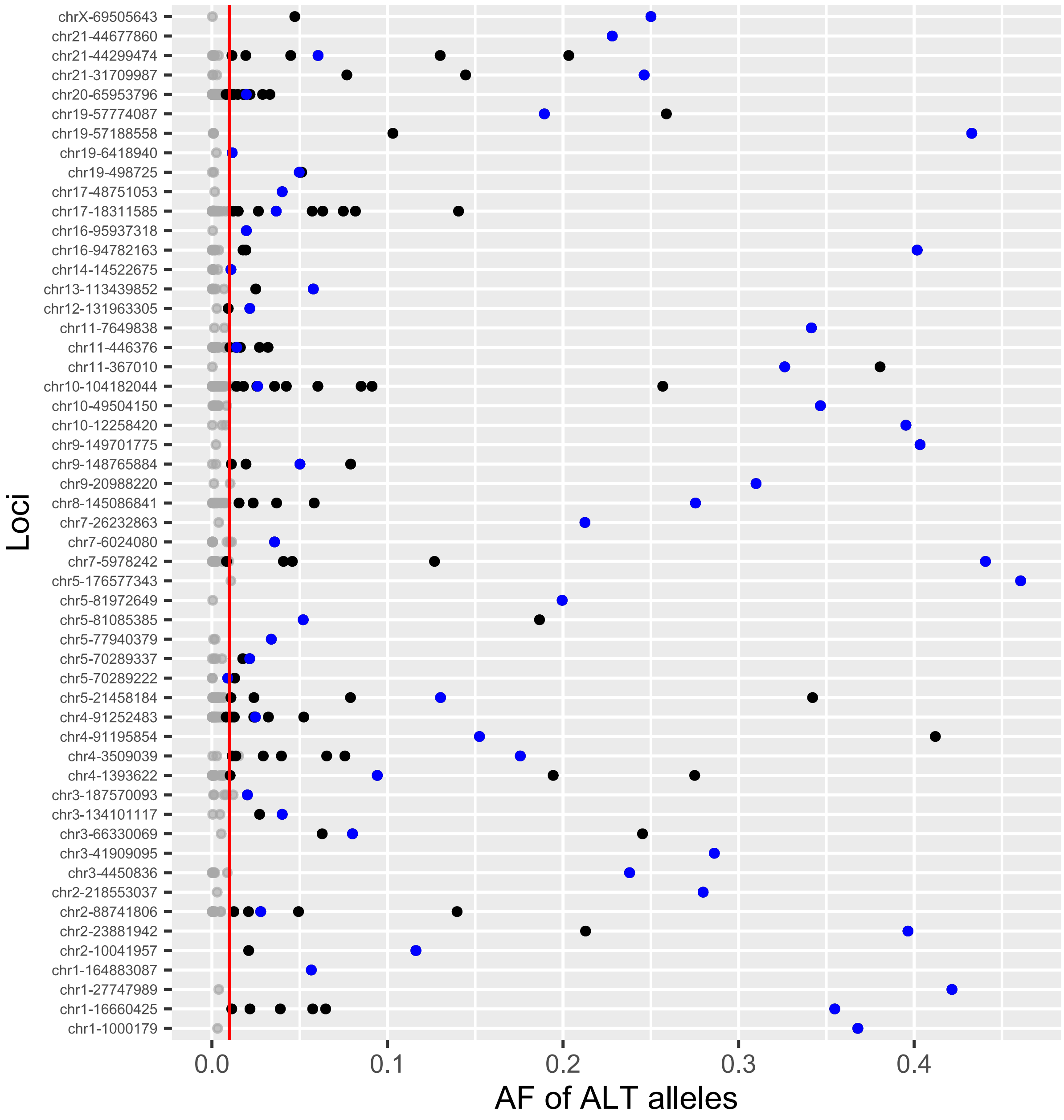

Loci
AF of ALT alleles
chrX-69505643
chr21-44677860
chr21-44299474
chr21-31709987
chr20-65953796
chr19-57774087
chr19-57188558
chr19-6418940
chr19-498725
chr17-48751053
chr17-18311585
chr16-95937318
chr16-94782163
chr14-14522675
chr13-113439852
chr12-131963305
chr11-7649838
chr11-446376
chr11-367010
chr10-104182044
chr10-49504150
chr10-12258420
chr9-149701775
chr9-148765884
chr9-20988220
chr8-145086841
chr7-26232863
chr7-6024080
chr7-5978242
chr5-176577343
chr5-81972649
chr5-81085385
chr5-77940379
chr5-70289337
chr5-70289222
chr5-21458184
chr4-91252483
chr4-91195854
chr4-3509039
chr4-1393622
chr3-187570093
chr3-134101117
chr3-66330069
chr3-41909095
chr3-4450836
chr2-218553037
chr2-88741806
chr2-23881942
chr2-10041957
chr1-164883087
chr1-27747989
chr1-16660425
chr1-1000179
0.0
0.1
0.2
0.3
0.4

**Supplementary Figure 9. *BORCS7* expression as a function of the cumulative number of repeats or the number of alternate insertion alleles at the multiallelic SV locus chr10:104182044.** (**A**) *BORCS7* expression (y-axis) is not associated with the cumulative number of “AC” motif repeats (x-axis) at chr10:104182044 in 504 CaG participants (P-value=0.81). Participants with 0 repeat do not carry the insertion. (**B**) We repeated the analysis in **A** but with participants that have 2 alternate (insertion) alleles (N=299). Again, the cumulative number of “AC” motif repeats is not associated with *BORCS7* expression (P-value=0.31). (**C**) The association between *BORCS7* and the number of insertion allele (0, 1 or 2; independently of the number of “AC” motif repeats) is also not significant (P-value=0.98). The blue dots represent the participants carrying one SV-eQTL allele (the allele associated with *BORCS7* expression). There are no participants carrying two copies of the SV-eQTL allele. The red lines denote the regression lines fitted to the data.

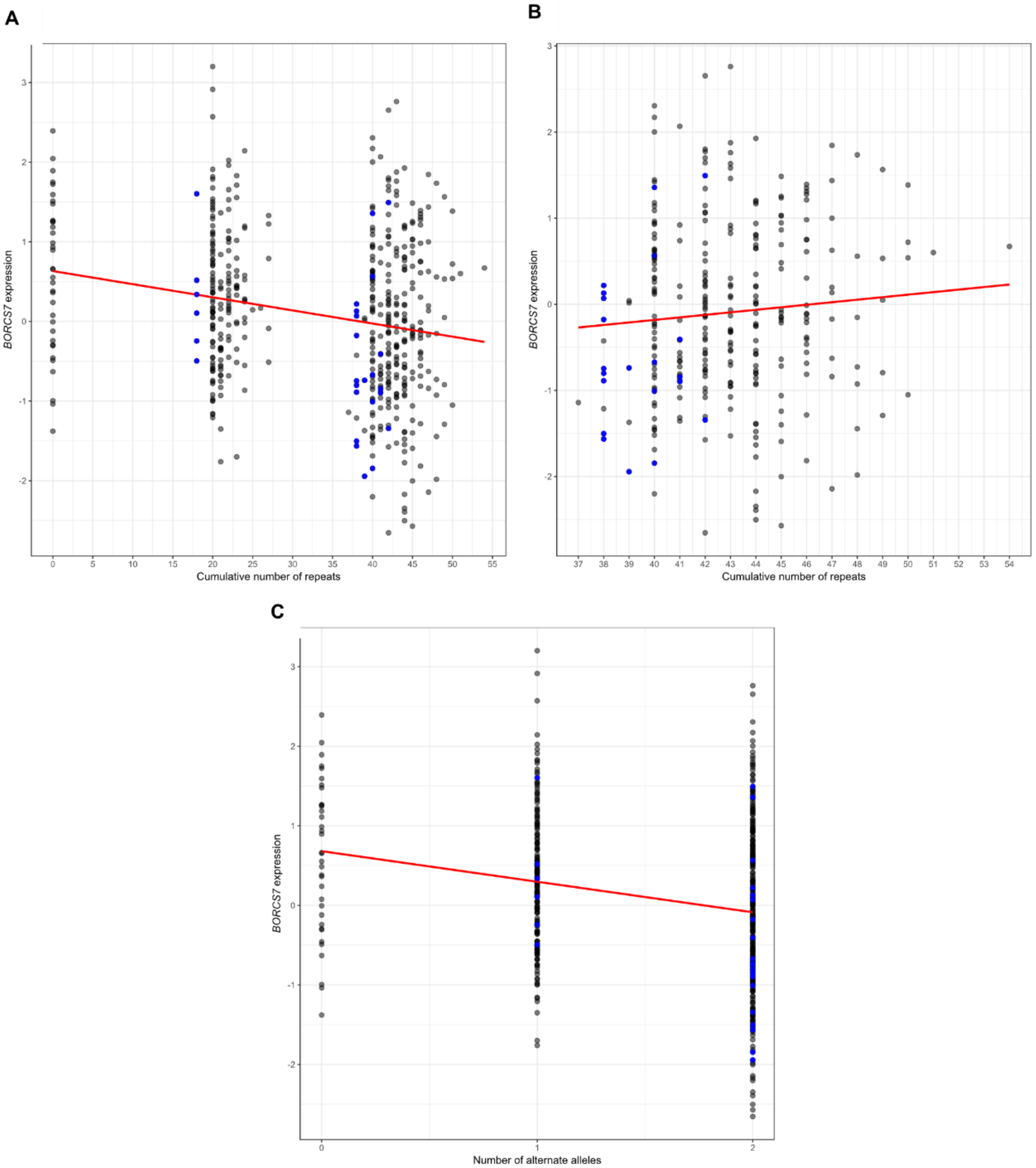
A
B
C
BORCS7 expression
Cumulative number of repeats
Number of alternate alleles
